# Human placental villi immune cells help maintain homeostasis *in utero*

**DOI:** 10.1101/2021.07.14.452362

**Authors:** Jessica M Toothaker, Oluwabunmi Olaloye, Blake T. McCourt, Collin C McCourt, Rebecca M Case, Peng Liu, Dean Yimlamai, George Tseng, Liza Konnikova

## Abstract

Maintenance of healthy pregnancy is reliant on successful balance between the fetal and maternal immune systems. Although maternal mechanisms responsible have been well studied, those used by the fetal immune system remain poorly understood. Using suspension mass cytometry and various imaging modalities, we report a complex immune system within the mid-gestation (17-23 weeks) human placental villi (PV). Further, we identified immunosuppressive signatures in innate immune cells and antigen presenting cells that potentially maintain immune homeostasis in utero. Consistent with recent reports in other fetal organs, T cells with memory phenotypes were detected within the PV tissue and vasculature. Moreover, we determined PV T cells could be activated to upregulate CD69 and proliferate after T cell receptor (TCR) stimulation and when exposed to maternal uterine antigens. Finally, we report that cytokine production by PV T cells is sensitive to TCR stimulation and varies between mid-gestation, preterm (26-35 weeks) and term deliveries (37-40 weeks). Collectively, we elucidated the complexity and functional maturity of fetal immune cells within the PV and highlighted their immunosuppressive potential.

## Introduction

Successful pregnancy is dependent on a delicate immune homeostasis, yet many of the factors required to maintain this homeostasis remain elusive. It is understood that the maternal immune system must balance pathogen defense while also preventing rejection of the semi-allogenic fetus for ~40 weeks in a healthy full term pregnancy (Erlebacher, 2013; PrabhuDas et al., 2015). Adding to the complexity, the progression of pregnancy is mirrored by distinct physiological states at many sites throughout the body requiring the maternal immune system to be equally dynamic and adaptive. This point is clearly illustrated in the seminal study by Agheepour and colleagues that tracked immunological responses in the periphery from early gestation through delivery using mass cytometry (CyTOF) (Aghaeepour et al., 2017). Many studies have identified numerous mechanisms by which the maternal immune system accommodates the developing fetus at the fetal-maternal interface. These include: the detection of suppressive uterine NK cells reviewed in (Gaynor and Colucci, 2017), presence of novel T regulatory (Treg) populations (Salvany-Celades et al., 2019), detection of suppressive B cells (Huang et al., 2017), restricted access to plasmacytoid dendritic cells (pDC) (Li et al., 2018) and a predominance of type 2 helper T cells (Miyazaki et al., 2003). The importance of the maternal immune system in pregnancy cannot be understated, however recent findings suggest that the fetal immune system must also be considered to fully understand placental immune homeostasis throughout gestation.

Historically, the fetal and neonatal immune systems were thought to be immature. This hypothesis was supported by poor vaccine responses in neonates (Saso and Kampmann, 2017), high susceptibility to infection (Simonsen et al., 2014), and the predominance of naïve lymphocytes in human cord blood (Paloczi, 1999). However, novel insights suggest that the fetal and neonatal immune systems are developed, though potentially have altered functions. Work supporting this concept includes: the *in utero* maturation following education of fetal Tregs (Mold et al., 2008), detection of novel immunosuppressive cell types present in neonatal blood (Elahi et al., 2013; Halkias et al., 2019; Miller et al., 2018a), and the presence of *in utero* memory lymphocytes in many fetal tissues (Li et al., 2019; Odorizzi et al., 2018; Schreurs et al., 2019; Stras et al., 2019a; Zhang et al., 2014). However, our knowledge about the immunological capabilities of the fetal cells at the fetal maternal interface is sparce.

A collection of recent single cell RNA-sequencing studies of the first trimester fetal-maternal interface revealed multiple previously undocumented PV cell types and cell-cell interactions (Suryawanshi et al., 2018; Vento-Tormo et al., 2018). Similarly, detection of novel cell populations and interactions was observed in third trimester placental surveys (Pavličev et al., 2017; Pique-Regi et al., 2019). Of interest, the work by Pique-Regi specifically identified PV-specific immune cell signatures, notably the presence of both resting and activated T cells of fetal origin in term PV (Pique-Regi et al., 2019). These data are in line with other work detecting a previously undocumented complex and diverse PV immune system in third trimester preterm rhesus macaques, which also contained T cells with activated phenotypes (Toothaker et al., 2020).

Building on these studies we hypothesized that the active PV immune system detected in the third trimester (Pique-Regi et al., 2019; Toothaker et al., 2020) must be present and contribute to immune homeostasis at mid-gestation. Using a combination of RNA-sequencing, CyTOF, imaging mass cytometry (IMC), and florescent microscopy, we investigated the PV immune profile from healthy mid-gestation (17-23 weeks) placental tissues. With this unique sample cohort, we detected multiple PV-specific immune signatures (absent in the decidua and membranes from the chorionic plate). We also identify that PD-L1 expression on antigen presenting cells is reduced in preterm placentas suggesting that PD-L1 expression may help keeps this armed PV immune system homeostatic *in utero.* Furthermore, using functional assays we uncovered that the PV T cells are poised to execute mature inflammatory functions as early as 18 weeks’ gestation, and that the cytokine secretion of PV T cells is variable in mid-gestation and preterm pregnancies but consistent across term pregnancies after T cell receptor (TCR) stimulation.

## Results

### The human mid-gestation PV have tissue specific immune signatures

We collected placental specimens from 19 second trimester products of conception, gestational age (GA) 17-23 weeks **(Table S1)**. Maternal decidua and fetal chorionic/amniotic membranes covering the chorionic plate (referred to hereafter as CP) were separated from the PV **(Fig 1A)** with forceps under a dissecting microscope. Separation of layers was initially confirmed by histology **(Fig 1B)**. Tissue was then either cryopreserved for CyTOF analysis, as previously described (Konnikova et al., 2018; Stras et al., 2019a) and validated in **Fig S1A**, fixed with formalin prior to embedding in paraffin for imaging mass cytometry (IMC) and immunofluorescence (IF) analysis or snap frozen for bulk RNA-sequencing (RNAseq). To verify separation of placental layers, we used bulk RNAseq from 3 matched cases (**Table S1**). Differential expression analysis (**Table S2**) and hierarchical clustering confirmed segregation of layers based on transcription profiles with the exception of one outlier (CP3) sample which was enriched for inflammatory signatures, likely upregulated during the D&E procedure or secondary to undocumented *in utero* inflammation **(Fig 1C)**. This segregation of samples was confirmed with k-means clustering which grouped samples correctly by tissue with the exception of CP3 outlier **(Fig 1C)**. Moreover, we confirmed the enrichment of decidua and PV specific stromal genes previously reported in multiple studies (Pique-Regi et al., 2019; Suryawanshi et al., 2018; Vento-Tormo et al., 2018) and detected two tubulin genes, TUPP3 and TUBAL3 enriched in all three CP samples. To determine if immune cells in the mid-gestation PV were solely reflective of the Hofbauer cell population we used immunofluorescence to co-stain for CD45, a marker of all hematopoietic cells and CD163, a classical PV resident Hofbauer cell marker (Reyes and Golos, 2018). Consistent with previous reports identifying non-Hofbauer immune subsets in the first and third trimester PV (Bonney et al., 2000; Pique-Regi et al., 2019), we detected CD45^pos^CD163^lo^ cells within the mid-gestation PV **(Fig 1D)** ranging in abundance from 30-70% of CD45^pos^ nuclei per high power field **(Fig S1B)**. As the PV are bathed in maternal blood (intervillous), we also confirmed that immune cells present in PV samples were reflective of cells contained within the trophoblast layers PV itself (intravillous) and not simply contamination from maternal blood cells **(Fig 1D, S1B)**. Additionally, we detected the Y chromosome with *in situ* hybridization in many intravillous cells and had enriched expression of Y chromosome derived *EIF1AY* mRNA coupled with low expression of the X chromosome inactivation transcript *XIST* in male PV samples indicating that the majority PV immune cells in our study were fetal in origin **(Fig 1E, S1C)**.To gain insight into what the CD45^pos^CD163^lo/neg^ immune cell subsets could be, we next assessed the expression of immune genes in our RNAseq dataset and confirmed the presence of a diverse immune landscape in mid-gestation placental tissues. Though most transcripts in the PV were expressed at lower levels than decidual counterparts (darker in color), transcripts for most major immune subtypes analyzed were detected in PV samples (circle size) including T cells, B cells, DCs and macrophages (Mϕs) **(Fig 1F, Table S3)**.

**Figure 1.**
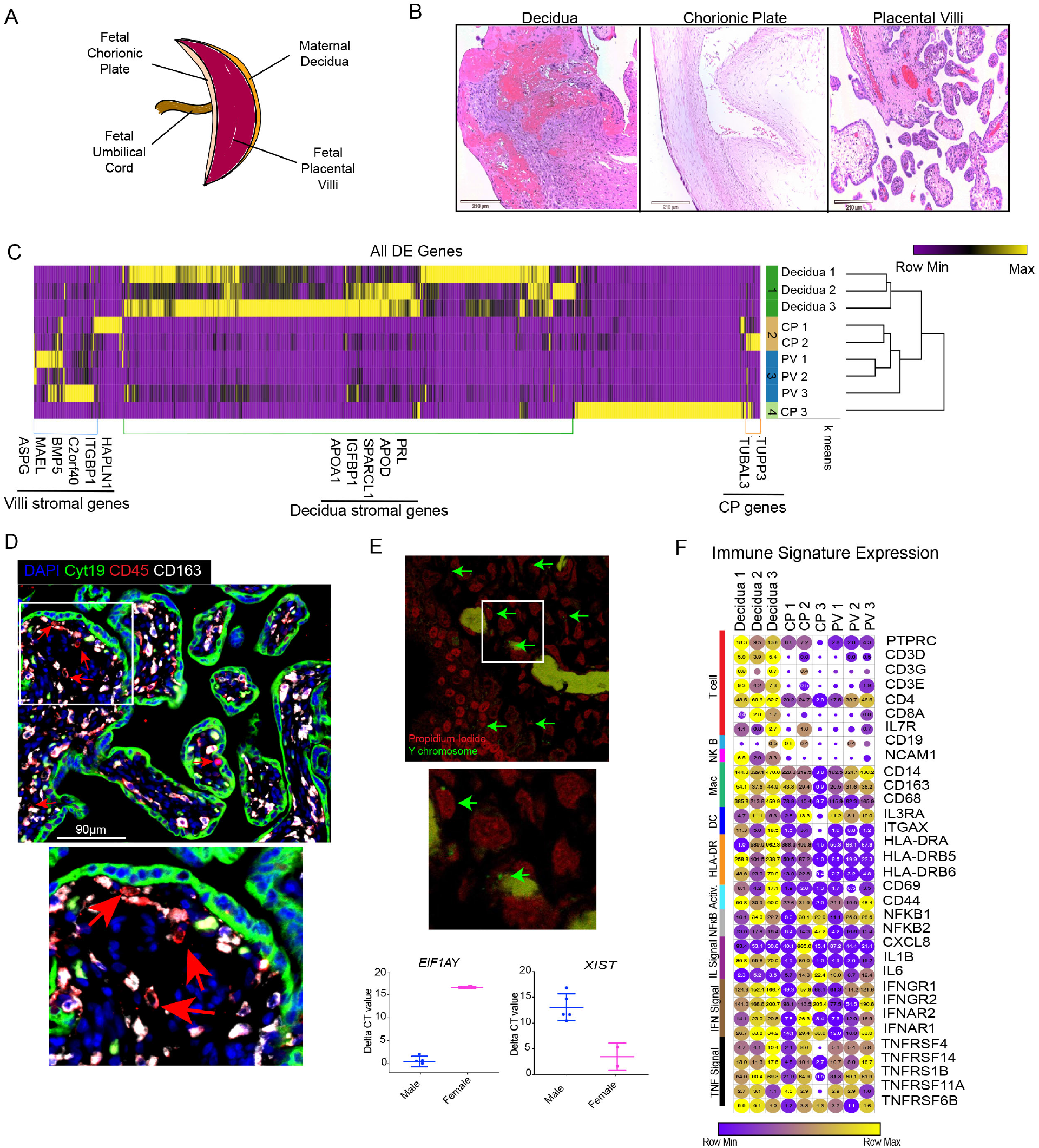
Tissue specific signatures in the mid-gestation placenta. (A) Diagram of placental tissues. (B) Representative H&E staining of layers of placental tissues. (C) All differentially expressed genes between three placental tissues with p value <0.05, false-discovery rate <20%, and fold change > absolute value 2. (D) Representative image of CD45^pos^ CD163^lo/neg^ cells within the intravillous space identified with immunofluorescence. (E) Representative image for fluorescent *in situ* hybridization of Y chromosome in PV (top). Delta CT values of Y and X chromosome genes in male and female PV (bottom). (F) Expression values of selected immune genes. Circle size indicative of expression value. Circle color indicative of relative expression across row. * = p value <0.05 upon post hoc analysis after Kruskal-Wallis (K-W) test. DE = differentially expressed. CP= chorionic plate.

To survey the CD45^pos^ populations in the PV, we used a panel of 38 metal conjugated antibodies **(Table S4)** and performed CyTOF analysis on 12 placenta-matched decidua, CP and PV samples **(Table S1)**. Briefly, cryopreserved tissues were then batch thawed and digested to make single cell suspensions, stained with metal conjugated antibodies (**Table S4)** and analyzed using CyTOF (Konnikova et al., 2018) **(Fig S1A)**. FCS files from CyTOF analysis were pre-gated for DNA^pos^, single, live, non-bead, CD45^pos^ cells **(Fig S2A)**. After omitting samples with insufficient cell numbers (>750 CD45^pos^ cells) we were left with 11 total samples for each tissue layer **(Table S5)**. CD45^pos^ cells were clustered using an automated clustering algorithm, Phenograph **(Fig 2A)** and clusters identified based on mean metal intensities of the surface markers from Clustergrammer generated associated heatmaps **(Table S6, Fig S2B)**.

**Figure 2.**
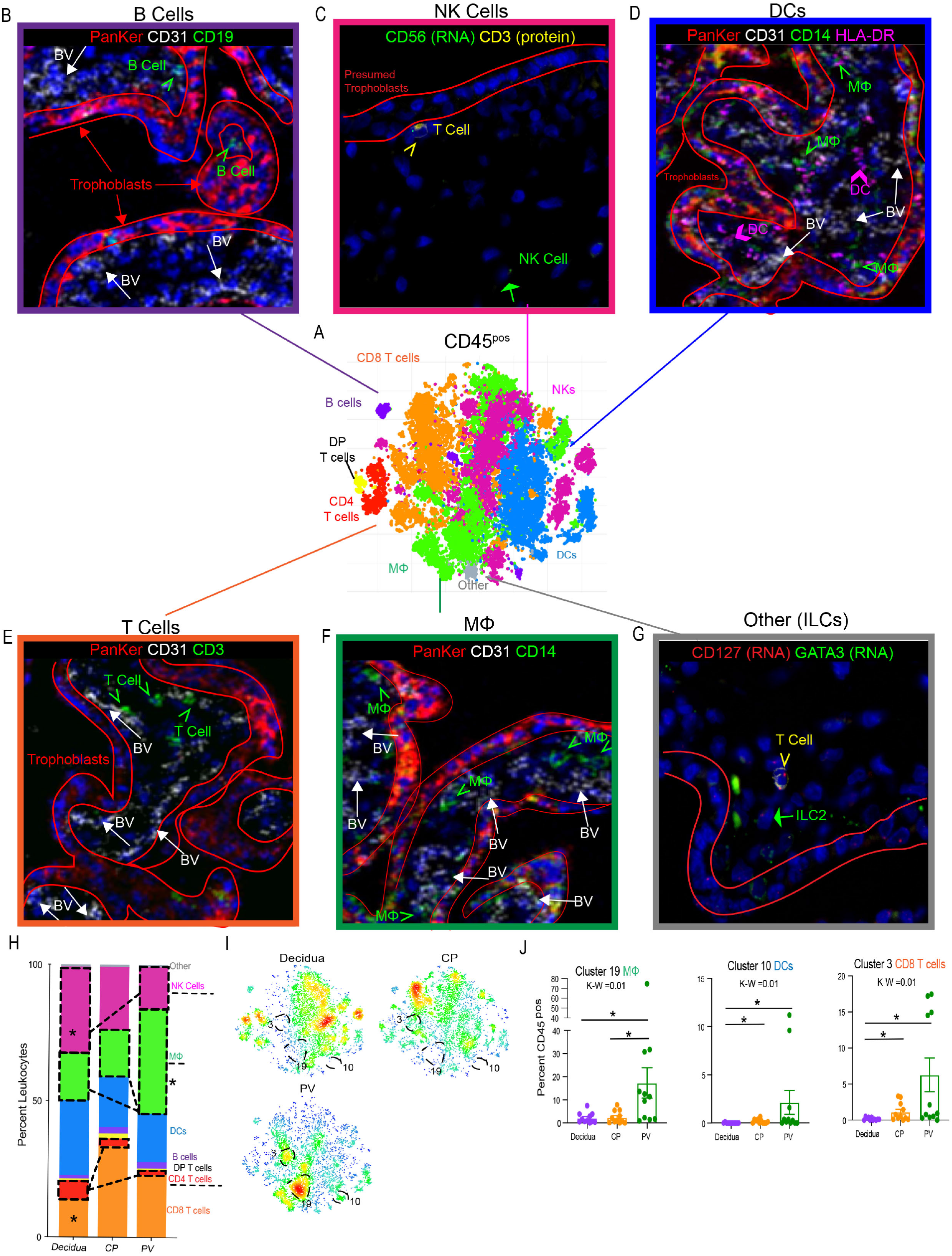
Global Immune Landscape of Second Trimester Placenta. (A) Merged tsne of CD45^pos^ cells from maternal decidua (n=11), CP (n=11) and PV (n=11) from Phenograph clustering of CyTOF data. (B) IMC identifying B cells (green) arrows located within the trophoblast bound (red) and outside the fetal blood vessels (BV) (white arrows). (C) Dual in situ hybridization and immunofluorescence (IF) identifying NK cells (green arrow) phenotypically distinct from T cell (yellow arrowhead) in PV. (D) IMC identifying DCs (pink arrows) and HLA-DR^pos^ Mfs (green arrows). (E) IMC identifying T cells (green arrows). (F) IMC identifying Mfs (green arrows). (G) Dual in situ hybridization and IF identifying ILC2s (green arrow) phenotypically distinct from T cell (yellow arrowhead) in PV. (H) Stacked bar graph comparing summation of all clusters belonging to same immune subsets from CyTOF data. (I) Density plot reflective of populations in (A) segregated by tissue of origin. Clusters significantly enriched in PV outlined. (J) Quantification of PV enriched cluster abundance. K-W = Kruskal-wallis test. * = p value <0.05 following posthoc analysis.

To confirm that PV immune subsets identified were not solely reflective of blood leukocytes in the fetal circulation, we used immunofluorescence (IF) and imaging mass cytometry (IMC) with a panel that included 23 markers **(Table S7)** on 6 total regions of 2 individual PV cases **(Table S1)**. Using IMC, we validated that B cells, DCs, T cells and Mϕs were found outside the fetal vasculature (CD31) in the PV stroma **(Fig 2B-F)**. Due to technical limitations in generating IMC antibodies to identify NK cells and ILCs, we used dual *in situ* hybridization and immunofluorescence to detect these populations in the PV as well **(Fig 2C,G)**. Though it is likely that some PV immune cells detected with CyTOF represent blood leukocytes, we can conclude that a proportion of the PV immune cells represent stromal populations.

To further characterize these cells, we used CyTOF analysis consisting of 31 unique clusters of immune cells within the STP, belonging to Mϕ, DC, NK, CD4 T cell, CD8 T cell, double positive (DP) T cell, B cell and other immune cell type subsets **(Fig 2H, S2B)**. Each layer of the placenta housed a unique and complex immune profile **(Fig S2C)**. When all clusters belonging to the same immune subset were combined, the decidua had a greater abundance of NK cells compared to PV **(Fig 2H),** consistent with previous studies (King et al., 1991; Koopman et al., 2003). Additionally, there was a higher proportion of CD4 T cells in the decidua than either of the fetal layers, likely attributed to the documented high abundance of decidual Tregs **(Fig 1H)**. In contrast, the PV had a larger proportion of Mϕ than either the decidua or CP, potentially representing the Hofbauer population **(Fig 2H)**.

When each cluster abundance was directly compared, 11/31 CD45^pos^ clusters were differently distributed between the three layers of the placenta **(Fig 2I-J, S2D)**. The PV was uniquely enriched for cluster 19 CCR7^neg^ Mϕs **(Fig 1I-J).** This robust cluster similarly suggests the presence of Hofbauer cells (as prior reports show most Hofbauer cells are CCR7^neg^ (Joerink et al., 2011)) in the PV and confirms the tissue-specificity of Hofbauer cells in our data set due to this cluster being largely undetectable in decidua and CP samples **(Fig 2J)**. Interestingly, we also found cluster 10, CCR7^neg^ DCs, and cluster 3, CD69^neg^ CD8 T cells, to be enriched in the PV over decidua **(Fig 2I-J)**. CCR7 is highly expressed on DCs homing to secondary lymphoid structures from peripheral tissues after antigen encounter (Ohl et al., 2004). CD69 is found to be transiently upregulated on activated T cells (Cibrián and Sánchez-Madrid, 2017) and constitutively upregulated on tissue resident memory T cells (Kumar et al., 2017). As PV enriched clusters 9 and 3 lacked these respective markers, we hypothesize that PV are poised (immune cell types are present) to execute mature immune function, such as antigen presentation, but may not be actively performing such functions during a time of homeostasis such as mid-gestation in healthy pregnancies. To further explore this idea, we next analyzed each immune cell subset more thoroughly.

### PV innate cells have quiescent phenotypes

To evaluate PV non-antigen presenting innate cell subsets independent of their antigen presentation abilities that represent the first cells to sense foreign antigens, we clustered on CD3^neg^CD19^neg^HLA-DR^neg^ cells (**Fig S3A)**. We identified Mϕs, innate lymphoid cells (ILCs), NK cells and multiple other immune cell populations that we were unable to fully phenotype with our panel **(Table S4), (Fig 3A, S3B)**. When comparing subtypes of immune cells, i.e. individual clusters of the same subtype added together, we confirmed our findings from the CD45 level (**Fig 1H)**, with the decidua having a larger proportion of NK cells and the PV having a larger proportion of Mϕs **(Fig 2B)**. The increased granularity of focusing on HLA-DR^neg^ innate cells specifically, revealed that Mϕs were both CD163^hi^ likely representing the Hofbauer cell population, and also contained a significant proportion CD163^lo^ Mϕs, presumably other non-Hofbauer cell Mϕs **(Fig 3B)**. HLA-DR^neg^ Mϕs in the decidua, in contrast were largely CD163^hi^, consistent with previous data on the phenotype of decidual Mϕs (Jiang and Wang, 2020) and with the enriched CD163 gene signatures seen in the decidua over CP from RNAseq data **(Fig 1D).** The Mϕ profile in the CP was more equally split between the two CD163^lo^phenotypes **(Fig 3B)**. At the individual cluster level, multiple clusters were enriched in either the decidua and/or CP **(Fig S3C)**. Within the PV there were enriched in cluster 25, CD163^lo^ Mϕs, and cluster 27, NK cells **(Fig 3C)**. While cluster 27 NK cells were abundant in all three placental layers and only slightly elevated in the PV, cluster 25 Mϕs were specific to the PV and were only minimally present in decidua and CP **(Fig 3C)**.

**Figure 3.**
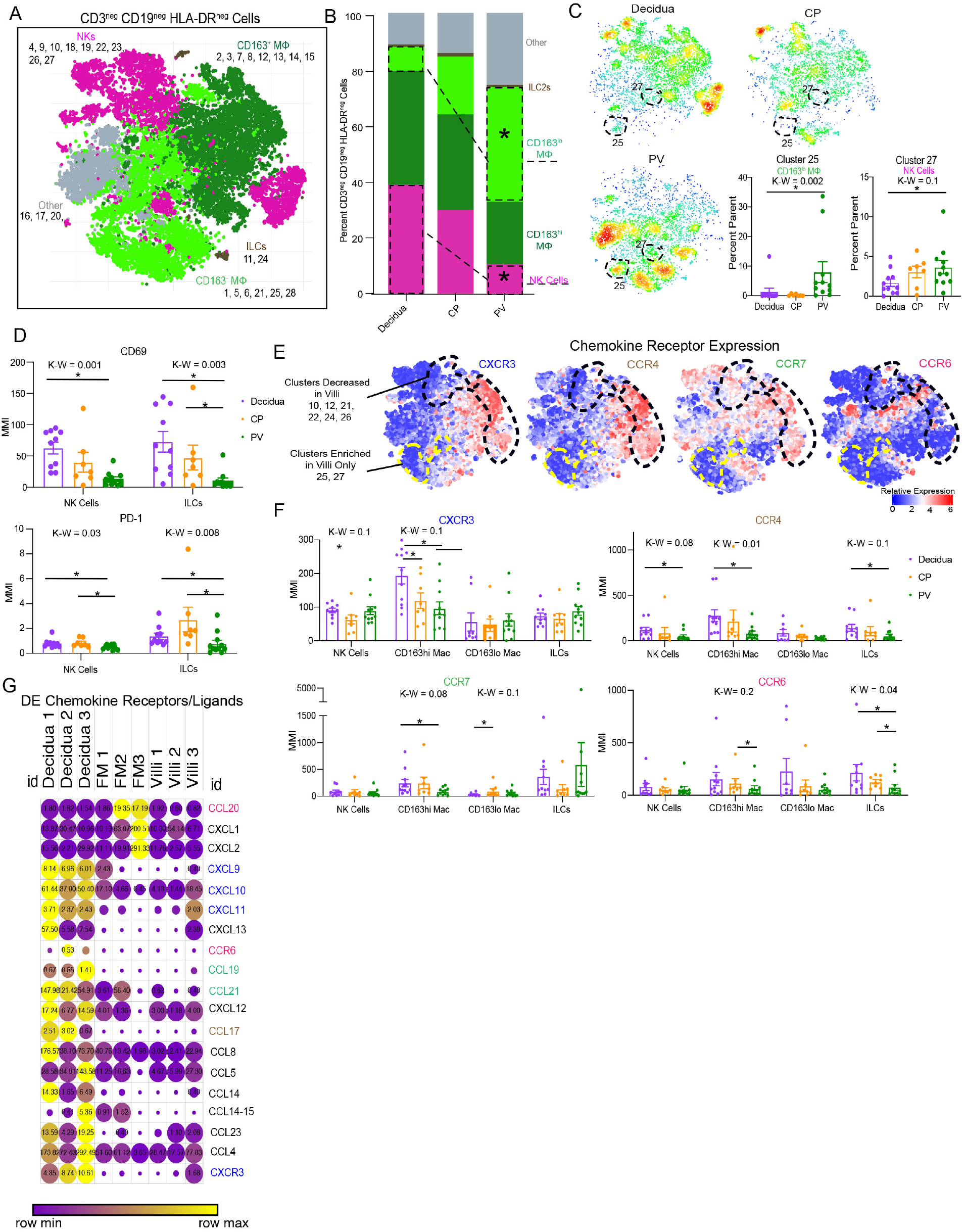
Innate HLA-DR^neg^ cells in PV. (A) Combined CyTOF tsne for CD45^pos^ CD3^neg^ CD19^neg^ HLA-DR^neg^ cells. (B) Stacked bar graph of abundance of major immune subtype. (C) Density plots separated by tissue of cell populations from (A). Statistically significantly abundant clusters in PV outlined. (right side) Cumulative data of PV abundant clusters outlined in density plots. (D) Mean metal intensities(MMI) of CD69 (top) and PD-1 (bottom) for 2D gated NK and ILC populations. (E) Expression heatmaps for chemokine receptors mapped to cells identified in (A). (F) MMIs of chemokine receptors on innate cell subsets from 2D gating. (G) Expression from RNA-sequencing of differentially expressed chemokine ligand/receptor genes between tissues. Circle size indicative of expression value, circle color reflective of relative expression across row. Differentially expressed determined as: p value <0.05, false-discovery rate <20%, and fold change > absolute value 2. * = p value <0.05 upon post hoc analysis after Kruskal-Wallis(K-W) test. DE= differentially expressed.

To determine if PV NK and ILCs were expressing markers consistent with activation, we compared the mean metal intensities (MMIs) of CD69 and PD-1 on these cells and found that PV NK cells and ILCs expressed significantly lower amounts of both CD69 and PD-1 than decidual and CP counterparts **(Fig 3D)**. Next, to examine if PV innate cells have migratory or tissue-retentive phenotypes we compared chemokine receptor (CCR) expression among subsets. Illustrated both visually **(Fig 3E)** and graphically **(Fig 3F)** we show that multiple populations of PV innate cells including NK cells, ILC and CD163^hi^ Mϕs had reduced expression of four CCRs. Interestingly, the expression of these markers between the three compartments was similar for the CD163^lo^ Mϕs (**Fig 3F**). To determine if other chemokine receptor/ligand pairs were also reduced in the PV, we identified 19 chemokine receptors/ligands that were differentially expressed between PV, CP and decidua using bulk RNA-seq **(Fig 3G, Table S8)**. These results validated the CyTOF findings of reduced expression of CCR6 and CXCR3 specifically, as well as at least one ligand for CCR7 and CCR4. Nine other chemokine ligand/receptors were implicated in this dataset **(Fig 3G).** These results suggest that PV innate cells are either static or are migrating in a non-signal specific manner and in combination with low expression of activation marker on PV immune cells, support a quiescent state of PV innate cells at mid-gestation.

### PV antigen presenting cells are diverse and phenotypically immunosuppressive

Next, we examined APC populations within each placental layer by clustering on CD45^pos^ CD3^neg^ HLA-DR^pos^ cells **(Fig S4A)**. We identified seven sub-types of APCs including: myeloid DCs (mDCs), plasmacytoid DCs (pDCs), CD163^hi^ Mϕs, CD163^lo^ Mϕs, B cells, NK cells and other cell types that we could not identify based on the available markers **(Fig 4A, S4B)**. In confirmation of our previous findings **(Fig 2-3)**, NK cells were again more prevalent in the decidua compared to the PV **(Fig 4B)**. HLA-DR^pos^ NK cells that are capable to independently present antigens to CD4 T cells have been described (Roncarolo et al., 1991). In contrast to HLA-DR^neg^ innate cells **(Fig 3)**, numerous individual APC clusters were enriched in the PV **(Fig 4C)**. Of note, multiple populations were enriched in the decidua and CP as well **(Fig S4C)**. Specifically, B cell clusters 7 and 27, mDC clusters 16 and 20, and CD163^lo^ CD4^neg^ Mϕ clusters 6 and 22 were significantly more abundant in the PV than either decidua or CP **(Fig 4C)**. Complimenting this finding, CD4^pos^CD163^hi^ Mϕs (cluster 26) was reduced in the PV compared to decidua and CP **(Fig S4C).** CD4^pos^ Mϕs have been shown to be long lived tissue resident Mϕs in the intestine and perhaps they serve a similar role in the decidua (Shaw et al., 2018). The large number of APC clusters (11 in total) differentially abundant between the PV and decidua/CP suggests that antigen presentation in the PV may be functioning through non-classical mechanisms at mid-gestation.

**Figure 4.**
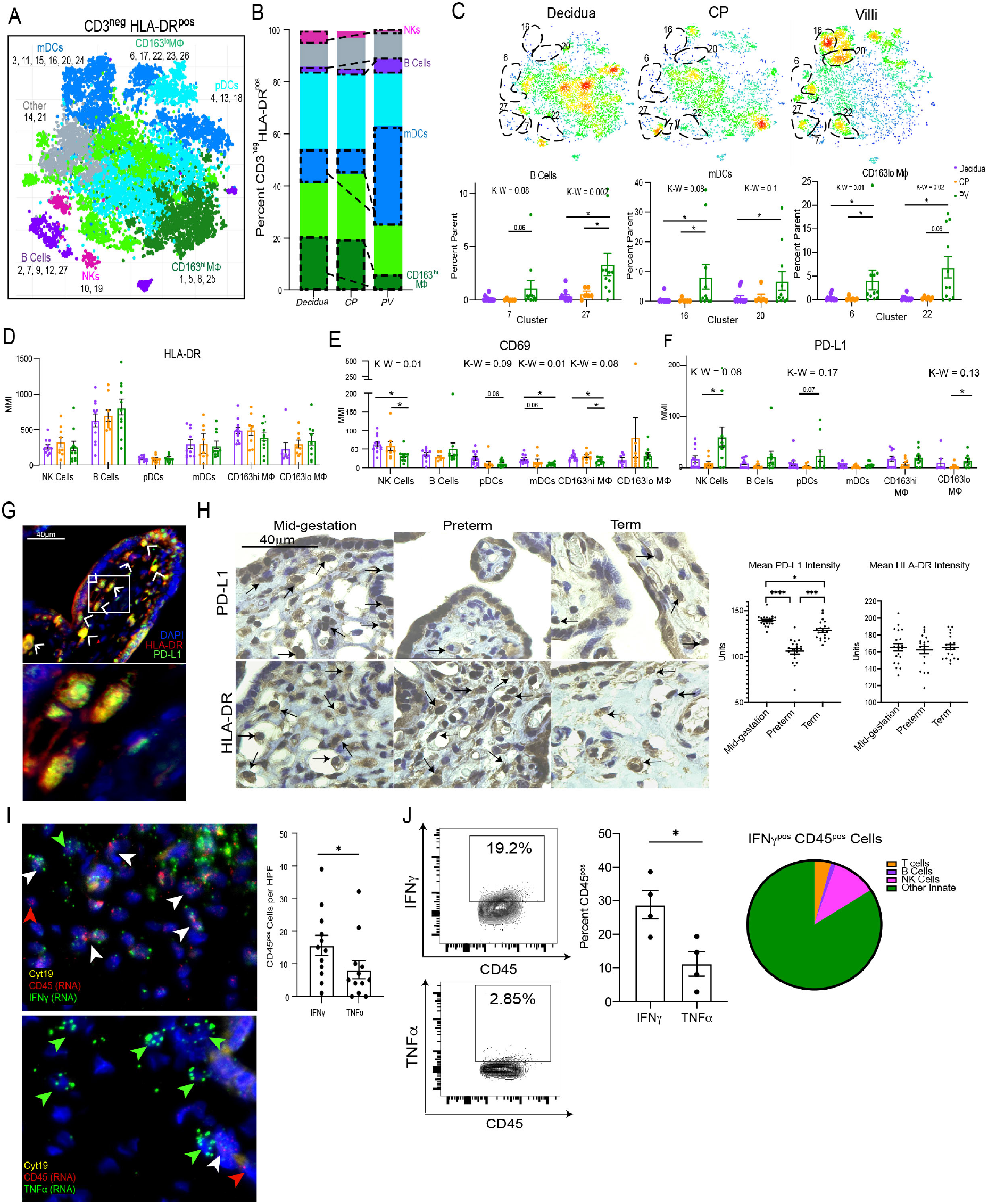
Antigen Presenting Cells in the PV. (A) Cumulative CyTOF tsne for CD45^pos^ CD3^neg^ HLA-DR^pos^ cells. (B) Stacked bar graph of abundance of major immune subtype. (C) Density plot separated by tissue of origin from (A). Statistically significantly abundant clusters in PV outlined. Cumulative data of PV abundant clusters outlined in density plots. MMI of HLA-DR (D), CD69 (E) and PD-L1 (F) for 2D gated populations. * = p value <0.05 upon post hoc analysis after Kruskal-Wallis test. (G) Representative image of PD-L1^pos^ APC populations in PV. (H) Representative images (left) and quantification of average stain intensity per stromal nuclei (right) for PD-L1 and HLA-DR IHC. (I) Representative images (left) and quantification of automated image analysis with CellProfiler (right) for dual RNA *in situ* hybridization and immunofluorescence in PV. (J) Representative flow plots and quantification for cytokine positive PV immune cells via flow cytometry. Pie-chart representing major immune subset abundance of IFNγ^pos^ immune cells from flow cytometry. * = p-value <0.05 in Mann-Whitney two-tailed test. K-W = Kruskal Wallis test p value. MMI = mean metal intensity.

To investigate potential functional distinctions in PV APCs, we examined the expression of both activation and immunosuppressive markers on each APC subset identified in **Figure 4B**. In contrast to the hypothesis that PV APCs have altered ability to function as APCs compared to decidua and CP, we found no difference in HLA-DR expression among APC subsets **(Fig 4D)**. However, consistent with PV APCs being more inhibited than decidua and CP counterparts, we identified significantly reduced expression of the activation maker CD69 in the PV CD163^hi^ Mϕs, NK cells, pDCs and mDCs **(Fig 4E)**. Furthermore, when we examined the inhibitory ligand, PD-L1, we documented its increased expression on multiple APC subsets, significantly so on CD163^lo^ Mϕs and HLA-DR^pos^ NK cells **(Fig 4F)**. This observation of high PD-L1 expression on PV APCs was confirmed by IF staining, where almost every observable PV HLA-DR^pos^ cell co-expressed PD-L1 **(Fig 4G, S4D)**.

To determine if PD-L1 expression on PV APCs is important for maintaining healthy pregnancy we compared both PD-L1 and HLA-DR expression in the PV stroma with immunohistochemistry (IHC) between healthy mid-gestation placentas (21-23 weeks’ gestation), preterm placentas from complicated pregnancies (29-35 weeks’ gestation) and healthy placentas delivered at full-term (39 weeks’ gestation) **(Fig 4H)**. Consistent with our CyTOF findings, mid-gestation stromal cells have high expression of PD-L1 per nuclei and PD-L1 staining patterns are congruent with stromal HLA-DR staining indicating that the PD-L1^pos^ cells analyzed were likely APCs **(Fig 4H)**. Of note, we discovered that stromal PD-L1 expression in preterm PV is significantly lower than that of mid-gestation and term PV. Furthermore, term PV also have significantly reduced PD-L1 expression on stromal cells than mid-gestation PV **(Fig 4H)**. In contrast, we observed increased PD-L1 expression on preterm trophoblasts compared to mid-gestation trophoblasts **(Fig S4E)**. To confirm that the reduction of stromal PD-L1 was not an artifact of reduced APC abundance in preterm and term placentas we compared mean HLA-DR expression between all three groups and found no differences **(Fig 4H)**. These results suggest PD-L1 expression on PV APCs helps maintain homeostasis during pregnancy illustrated by the highest expression during the mid-gestation window and significantly reduced expression in complicated (preterm) pregnancies.

To investigate the regulation of constitutive PD-L1 expression by PV APCs we measured the expression of IFNγ, a well-documented regulator of PD-L1 (Garcia-Diaz et al., 2017). We hypothesized constitutive expression of PD-L1 on PV APCs may be resultant from IFNγ (a regulator of PD-L1) production. Consistent with this hypothesis we found preferential transcription of IFNγ over TNFα at baseline by PV immune cells **(Fig 4I, S4F).** Additionally, PV immune cells transcribe comparable IFNγ, but transcribe decreased TNFα than decidual immune cells **(Fig S4G)**. The preferential production of IFNγ to TNFa was confirmed in PV immune cells with flow cytometry. We report this IFNγ was mostly derived from non-NK cell innate immune cells **(Fig 4J)**. As such, it is possible that PV immune cells produce IFNγ during homeostasis to drive expression of PD-L1 on APCs.

### The second trimester placenta is dominated by memory CD8 T cells

As we observed high PD-L1 expression on APCs, we next explored if there were T cells present in the PV that would be inhibited by immunosuppressive APCs. We identified both circulating (within blood vessel) and potentially tissue resident T cells in the PV **(Fig 5A, S5A)**. Moreover, using IMC, we surprisingly identified non-circulating T cells of CD4, CD8 and double negative (DN) phenotypes that expressed CD45RO, a marker upregulated after antigen experience and absent on naïve T cells **(Fig 5B, S5B)**. Turning to our CyTOF data, when clustering specifically on T cells **(Fig S5C)**, we found that all three layers of the placenta had a T cell profiles dominated by CD8 T Cells **(Fig 5C-D, S5D)**. Building off the initial detection of CD45RO by IMC, we found that the majority of T cells in the PV were of memory phenotypes, delineated based on expression of CCR7 and CD45RA **(Fig 5E)**. Additionally, we found CD8 T cells with tissue-resident memory (TRM) phenotype in all three layers **(Fig 5E)**. We detected CD69^pos^ T cells both as a marker of TRMs on CCR7^neg^ CD45RA^neg^ T cells and also among other T cell subsets **(Fig 5E)**. The expression of CD69 on multiple T cell populations strongly suggests that some populations of PV T cells are stromal and not reflective of fetal blood T cells, because recent work has shown that blood CD8 T cells do not express CD69 (Buggert et al., 2020). The detection of both CD69^neg^ and CD69^pos^ CD8 T cells in the PV is consistent with the enrichment of one cluster of CD69^neg^ CD8 T cells from our initial CD45^pos^ clustering **(Fig 2)**.CD8 and CD4 non-Treg cell subtypes were evenly distributed between all three layers **(Fig 5D)**. However, CD4 Tregs were enriched in the decidua compared to the PV **(Fig 5D)**. The abundance and importance of Tregs throughout pregnancy in the decidua is well documented (Mjösberg et al., 2010; Salvany-Celades et al., 2019), but the role of Tregs in the CP and the PV is unclear, though we have shown PV Tregs function abnormally during intraamniotic inflammation (Toothaker et al., 2020). Moreover, there was an enrichment of CD4^neg^ CD8^neg^ that were also CD56^neg^ and CD16^neg^ double negative (DN) T cells in the PV **(Fig 5D, S5D)**. These DN T cells were likely γδ T cells whose presence in the first trimester PV has been described (Bonney et al., 2000).

**Figure 5.**
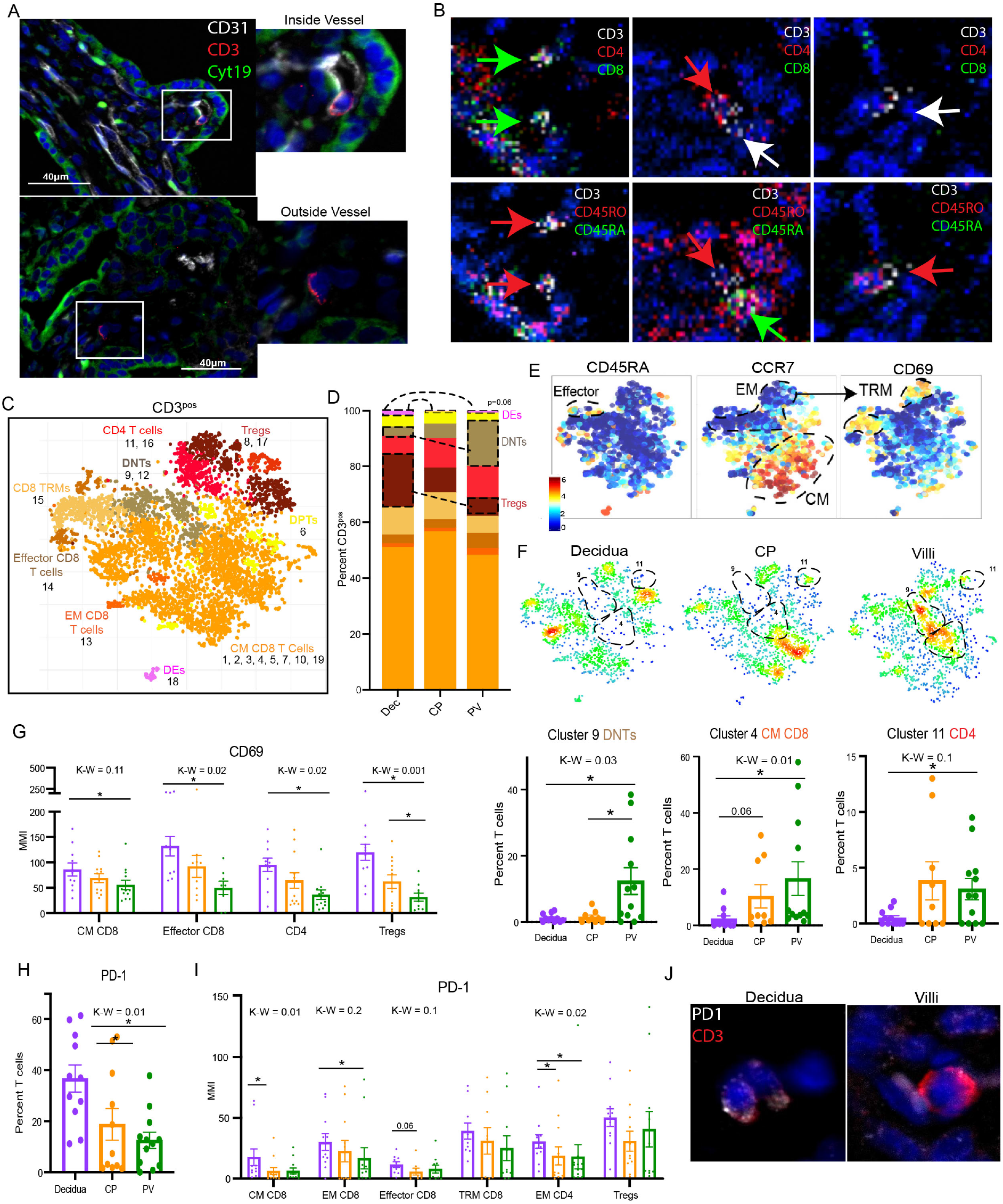
T cell subsets in placental tissues. Representative images of T cells inside (top) and outside (bottom) fetal vasculature (CD31) in PV. (B) IMC images of T cell subtypes in PV. (C) Cumulative CyTOF tsne for PV, CP and decidua CD45^pos^ CD3^neg^ HLA-DR^pos^ cells. (D) Stacked bar graph of abundance of major immune subtype. (E) Relative expression of memory T cell markers in PV cell populations from (C). (F) Density plot separated by tissue of origin (C). Statistically significantly abundant clusters in PV outlined. Cumulative data of PV abundant clusters outlined in density plots. (G) MMI of CD69 from 2D gating of populations. (H) Abundance of PD-1^pos^ T cells by 2-D gating. (I) PD-1 MMI from 2D gated subsets. (J) Representative image of PD-1 on T cells in PV and decidua. * = p value <0.05 upon post hoc analysis after Kruskal-Wallis (K-W) test. MMI = mean metal intensity. CM = central memory, EM = effector memory, TRM = tissue-resident memory.

The detection of PV T cells expressing memory markers was recently reported in the third trimester rhesus macaque (Toothaker et al., 2020). However, their identification in human mid-gestation PV is novel. To investigate T cell signatures unique to the PV, we next compared the abundance of individual T cell clusters. While two clusters were enriched in the decidua **(Fig S5E)**, cluster 9 CD4^neg^ CD8^neg^ T cells, cluster 4 CM CD8 T cells, and cluster 11 CD4 T cells were enriched in the PV **(Fig 5F)**. Cluster 11 T cells were CCR4^pos^CXCR3^neg^CCR6^neg^ **(Fig 5F, S5D)**, surface marker expression pattern suggestive of a TH2 phenotype, however further analysis for detection of TH2 specific transcription factors (GATA3) is needed for confirmation.

### Resting signatures define PV T cell subsets

Since there was a resting, homeostatic trend in PV innate cells **(Fig 2)** and a resting/coinhibitory profile among PV APCs **(Fig 4)**, we next explored if a resting phenotype was consistent among PV T cell subsets identified in **Fig 5D**. Among non-TRM T cells, PV T cell subsets exhibit reduced expression of CD69 **(Fig 5G)**. Moreover, the PV housed fewer PD-1^pos^ cells **(Fig 5H)** and reduced PD-1 per T cell compared to decidual counterparts **(Fig 5I-J)**. Though PD-1 is a marker of T cell exhaustion, it is also upregulated upon activation of the T cell receptor (summarized in (Xu-Monette et al., 2017)). We propose this is the more likely role of observed down-regulation of PD-1 in PV T cells as it is consistent with the downregulation of CD69.

### Maternal antigens can activate PV T cells

To determine if PV T cells have a reduced activation profile due to functional abnormalities or immaturity, we scanned for the presence of activated T cells via the detection of HLA-DR, phosphorylated Histone H3, phosphorylated S6 and phosphorylated CREB in T cells using IMC. Based on the above markers we were able to identify both resting **(Fig 6A)** and activated T cells in the PV **(Fig 6B)**. These results suggest that PV T cells are capable of being activated, consistent with our previous study in non-human primate which demonstrated TCR activation during *in utero* inflammation (Toothaker et al., 2020). As such we questioned if mid-gestation PV T cells could be activated in a TCR dependent manner. To test the functionality of the TCR pathway in PV T cells, we stimulated single cell suspensions from PV with soluble αCD3 and αCD28 antibodies for 72 hours. Consistent with normal TCR pathway functionality, we observed increased CD69 expression after stimulation **(Fig 6C)** and an increased proportion of cells proliferating **(Fig 6D, S6B-C)** with stimulation. Validating our previous findings that PV T cells are fetal in origin and unique from the decidua and other fetal organs, we observed an increase in abundance of rapidly proliferating T cells identified by high Ki67 expression (Miller et al., 2018b) after 4 hours of stimulation with αCD3 and αCD28 antibodies in fetal T cells (placenta and intestine) compared to adult T cells (decidua and intestine) **(Fig 6E, S6D)**. Of note, the placenta had the highest proportion of rapidly proliferating cells across all organs and ages following stimulation **(Fig 6)**.

**Figure 6.**
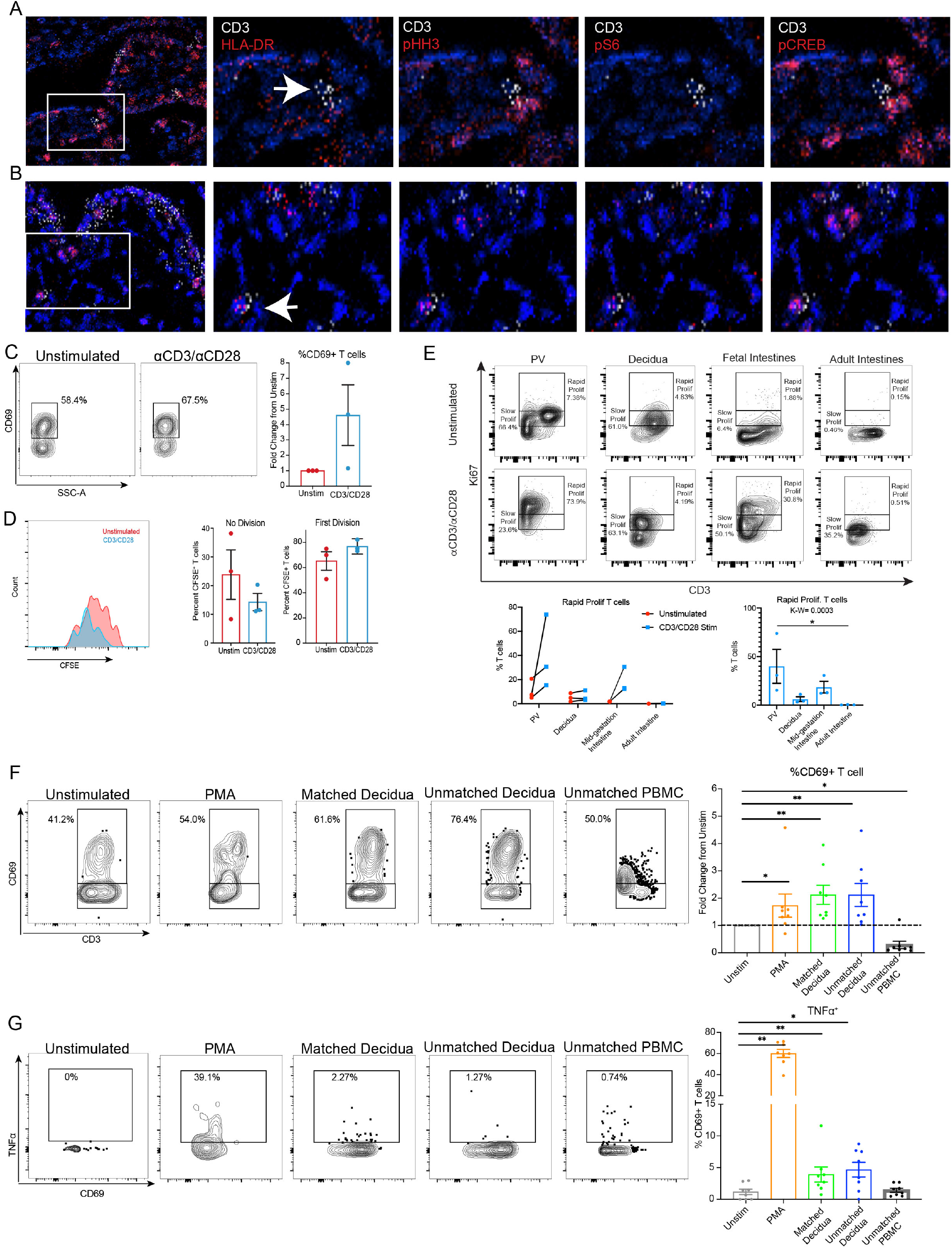
T cell activation in PV. (A) IMC images of inactive T cell in PV. (B) IMC images of activated T cell in PV. (C) Representative flow plots (left) and quantification (right) of CD69^pos^ T cell population. (D) Histogram showing proliferation as tracked by CFSE dye (left). Quantification of cells in proliferative subsets identified. (E) Representative flow plots (top) and quantification (bottom) for proliferating (Ki67^hi^) T cells after stimulation across tissues indicated. (F) Representative flow plots (left) and quantification (right) of CD69^pos^ T cell population. (F) Representative flow plots (left) and quantification (right) of TNFa^pos^ CD69^pos^ T cell population. * = p-value <0.05 in Mann-Whitney test.

To discern what antigens could activate PV T cells, we stimulated isolated PV cells with either PMA/Ionomycin as a positive control or lysed cellular components from either: pregnancy-matched decidua, unmatched decidua or unmatched pooled donor PBMCs. Consistent with activation of PV T cells, we saw elevated CD69 expression in the PMA/Ionomcyin, matched and unmatched decidual components conditions and but not with unmatched PBMC components **(Fig 6F**), suggesting that some antigens present in the decidua are capable of activating PV T cells. Validating the activation of PV T cells by decidual antigens, we also observed increased production of TNFα in a significant (though minor) proportion of CD69^pos^ T cells **(Fig 6G, S6E-F)**. These functional assays show that PV T cells have the potential to be activated by PV APCs through the TCR pathway and may become proinflammatory when stimulated in a TCR-dependent manner when exposed to particular antigens present in the uterine environment.

### Cytokine profile of PV T cells differentiates between mid-gestation, preterm, and term pregnancies

As we observed activation potential for mid-gestation PV T cells **(Fig 6)**, we next explored if PV T cell activation was consistent across gestation and between healthy and complicated pregnancies. To assess T cell activation at baseline in the absence of any ex-vivo T cell activation as consequence to tissue digestion and cell isolation, we measured phosphorylated (p)ZAP70 expression with IHC. ZAP70 is phosphorylated following engagement of the TCR. Interestingly, we observed increased pZAP70 intensity per ZAP70^pos^ stromal cell with increasing gestation (Term >preterm >mid-gestation) **(Fig 7A)**. Building upon this observation that TCR activation is variable between groups, we next assessed T cell function following a 4-hour stimulation with αCD3 and αCD28 antibodies. In confirmation of our CyTOF findings **(Fig 6)**, we report that the majority of PV T cells are CD8 T cells **(Fig S7A-C)**. We observed activation of CD8^pos^ T cells and very little activation of CD8^neg^ T cells via CD69 expression following stimulation in all three groups **(Fig 7B, S7A-B)**. We therefore chose to analyze activated (CD69^pos^) CD8 T cells in subsequent analysis. Surprisingly, we found that the production of four cytokines, TNFα, IL-17A, GranzymeB (GRZB), IFNγ, and did not vary between the three groups at baseline **(Fig 7C-F)**. Interestingly, the potential for CD4 PV T cells to make GRZB has been reported in the non-human primate PV (Toothaker et al., 2020). Consistent with this finding we identified high production with of GRZB from CD8 CD69^pos/neg^ T cells, CD8^neg^ T cells and NK cells **(Fig S7D)**. This finding Perhaps indicates an intrinsic function of PV T cells and NK cells as a second layer of antipathogen protection if the trophoblast layer is breached. In contrast to GRZB **(Fig 7E)** production of IL-17A, IFNγ, and TNFα was observed by ~30-60% of CD69^pos^ CD8 T cells **(Fig 7C-D,F)**.

**Figure 7.**
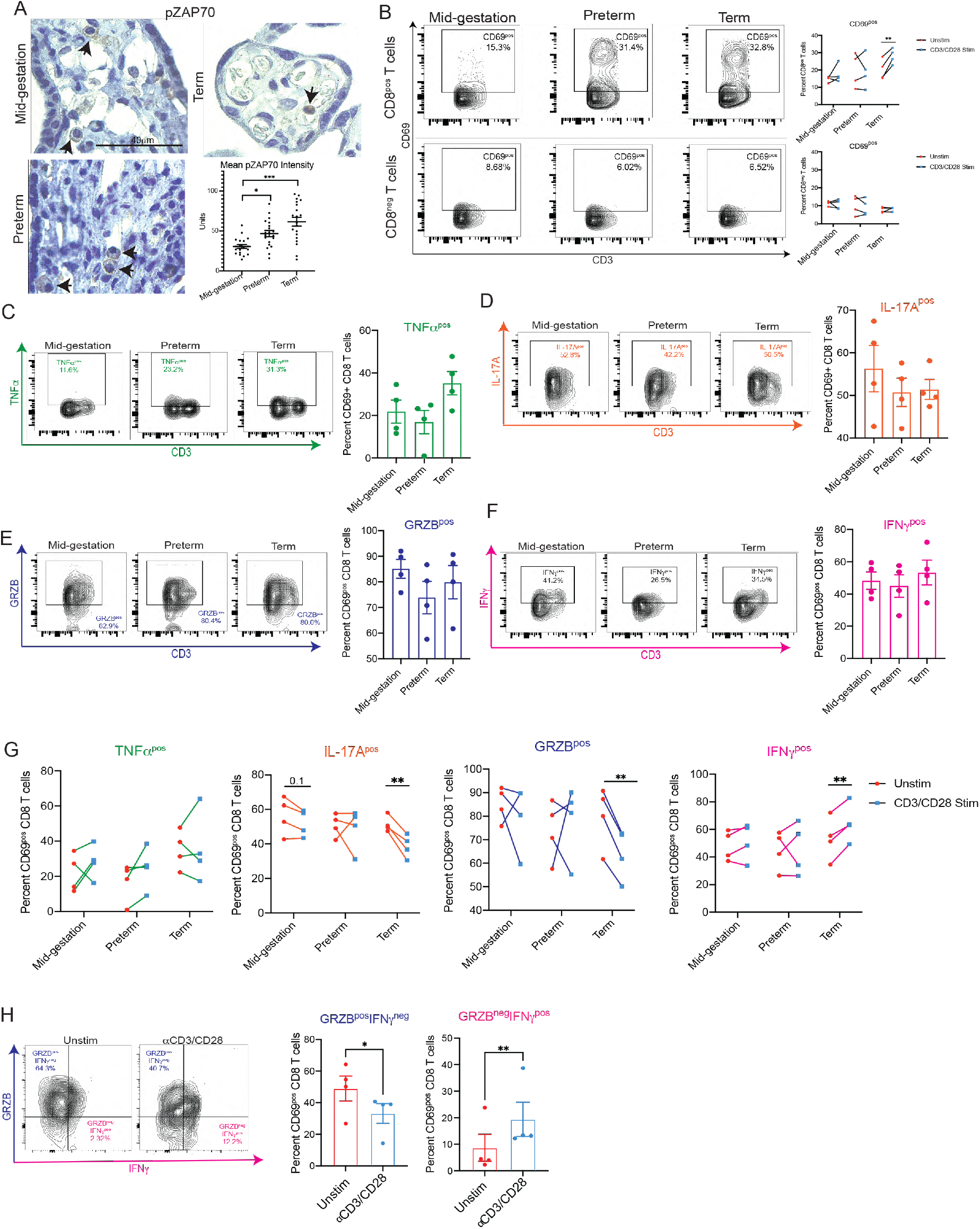
Gestational variation in PV T cell response. (A) Representative images and quantification of pZAP70 expression via IHC. (B) Representative flow plots and quantification of CD69 expression in CD8^pos^ and CD8^neg^ T cells following stimulation. (C-F) Representative flow plots and quantification of TNFa (C), IL-17A (D), GranzymeB (GRZB) (E), and IFNγ (F) at baseline. (G) Quantification of cytokine production before and after stimulation by CD69^pos^ CD8^pos^ T cells* = p-value <0.05 in Mann-Whitney test. (H) Representative flow plots of GRZB and IFNγ production by CD69^pos^ CD8^pos^ T cells* = p-value <0.05 in Mann-Whitney test.

Although baseline production of cytokines was consistent between mid-gestation, preterm, and term PV T cells, alterations in cytokine production occurred following stimulation. Across all four cytokines PV T cells isolated four distinct term placentas responded consistently. In contrast, mid-gestation and preterm PV T cells had high patient-to-patient variability **(Fig 7G)**. When tracking cytokine production of term PV T cells specifically, we observed no change in TNFα production. Additionally, we observed significant downregulation of IL-17A and GRZB following stimulation complimented with an increase in IFNγ production **(Fig 7G)**.To determine if the CD69^pos^ CD8 T cells were downregulating GRZB in favor of IFNγ we gated the two cytokines against one another **(Fig 7H).** Consistent with skewing of cytokine production from GRZB to IFNγ, there was a significant decrease of GRZB single positive cells and a concurrent increase of IFNγ single positive cells with stimulation **(Fig 7H)**. No significant differences were observed in either double positive or double negative populations **(Fig S7E)**. Importantly, this alteration from GRZB to IFNγ production was specifically observed in CD69^pos^ CD8 T cells and was not observed in other T cell subsets **(Fig S7F)**. Collectively, this experiment revealed that CD69^pos^ CD8 T cells in the PV are capable of producing a variety of cytokines at baseline and in full term healthy pregnancies a unique subset of CD69^pos^ CD8 T cells skews its cytokine production to favor IFNγ production upon stimulation of the TCR pathway.

In summation **(Fig 8)**, we demonstrate that the intravillous compartment of the healthy second trimester PV contains a diverse immune landscape comprised of innate cells such as Mϕs, NK cells and ILCs, as well as APCs including B cells, and memory T cells **(Fig 2)**. Moreover, we determined that the immune cells in the PV are likely capable of eliciting an inflammatory response, but instead maintain immune homeostasis at baseline through a variety of mechanisms. The mechanisms described include reducing chemotaxis by reduced expression of chemokine receptors on innate cells and low transcription of chemokine ligands in the PV overall **(Fig 3)**. Moreover, PV APCs constitutively express PD-L1, possibly regulated by the high expression of IFNγ by PV immune cells **(Fig 4)**. This PD-L1 expression by APCs is potentially needed to prevent the activation of antigen-experienced PV T cells **(Fig 5)** which can be activated through the TCR pathway by antigens present in the uterine environment **(Fig 6)** and result in the secretion of a variety of cytokines **(Fig 7)**.

**Figure 8.**
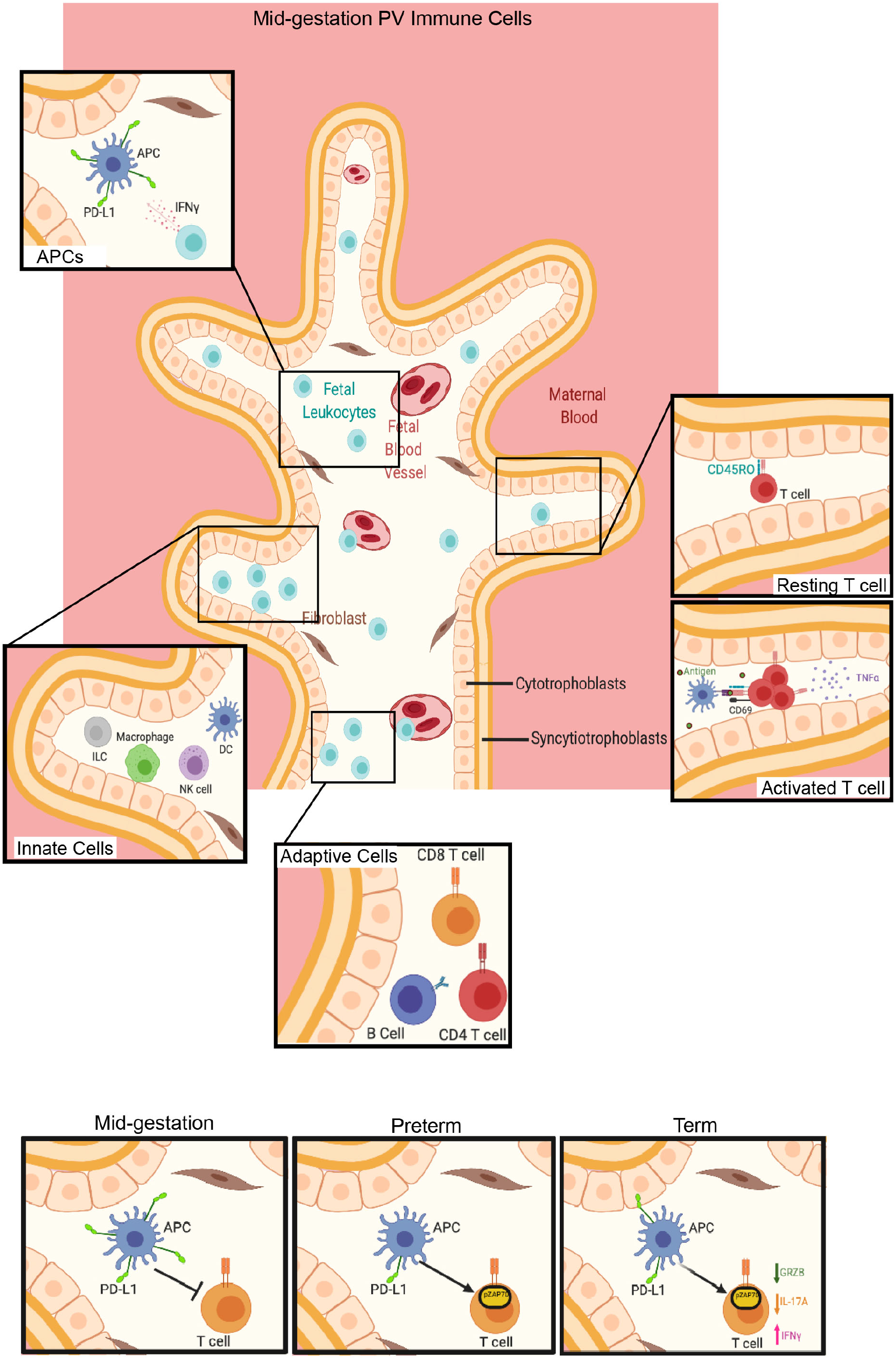
Summary of major findings in study. Image was generated with Biorender.com.

## Discussion

Preserving tolerance at the fetal-maternal interface is critical for maintaining a healthy pregnancy. Studies of the roles of maternal immunity within the decidua and maternal peripheral blood have resulted in the discovery of important immunological tolerance mechanisms including: tolerogenic uterine NK cells (Erlebacher, 2013), unique populations of Tregs (Salvany-Celades et al., 2019) restriction of DC migration to the uterus (Tagliani and Erlebacher, 2011) and suppressive B cells (Huang et al., 2017). Until recently however, the contribution of leukocytes within the PV beyond Hofbauer cells (PV resident Mϕs) has been far less explored.

Single cell studies within the past three years of the first trimester (Suryawanshi et al., 2018; Vento-Tormo et al., 2018) and full term (Pavličev et al., 2017; Pique-Regi et al., 2019) placenta highlighted the diversity of immune cells within the PV. Interestingly Pique-Rige and colleagues identified leukocyte signatures specific to the PV and highlighted a potential role for activated PV T cells in post-delivery PV (Pique-Regi et al., 2019). A finding also reported in preterm macaque PV (Toothaker et al., 2020). With these first and third trimester studies in mind, we hypothesized that the activated leukocytes detected by Pique-Regi (Pique-Regi et al., 2019) in humans and our group in non-human primates (Toothaker et al., 2020) may be present and poised to be activated in the PV earlier in gestation and subject to immunosuppressive mechanisms which maintain immune homeostasis within the PV in utero.

To answer these questions, we analyzed placental tissue (including decidua, PV and CP) and confirmed that there are diverse immune cells in second trimester PV samples that are located within the intravillous space and largely fetal in origin as suggested by the presence of the Y chromosome in majority of intravillous cells via FISH and the enrichment of Y genes in male fetuses. Although it is still possible that our analysis of PV immune cells included a proportion of maternal cells, ethical limitations precluded us from including maternal peripheral blood in our study. Future studies employing dual HLA-haplotyping/Y chromosome detection with staining for various immune populations on the same PV sections is needed to distinctly determine the fetal versus maternal origin of individual immune cell populations within the PV vasculature and stroma.

Previous studies have suggested that PV fetal immune cells are limited to Hofbauer cells (Thomas et al., 2021) However, the increased granularity provided by single cell methods, have allowed for the detection of T cells, B cells and NK cells of fetal origin in healthy term placentas (Pique-Regi et al., 2019). Moreover, a recent study identified infiltrating cells in the PV to be largely of fetal origin in cases of infectious villitis (Enninga et al., 2020) and Erbach et *al.* isolated T cells from single cell suspension at 18-24 weeks’ gestation (Erbach et al., 1993). It is important to note that studies using single cell suspensions make the delineation of immune cells from the PV vasculature indistinguishable from those in the PV stroma. In the current study, by combining CyTOF to capture the diversity of PV immune cells and multiple imaging modalities, we identified a diverse immune landscape in the PV that was distinct from both the decidua and CP populations consisting of both innate and adaptive immune cells, including T cells, present outside the fetal vasculature in the intravillous space.

Focusing specifically on the innate non-APC population we identified NK cells, ILCs and Mϕs to be present in the PV. Thomas et *al.* recently reported that Hofbauer cells in the first trimester are HLA-DR^neg^ (Thomas et al., 2021), consistent with this we found an increased abundance of Mϕs in the PV compared to the decidua in the innate HLA-DR^neg^ compartment. Surprisingly, we report that many of these HLA-DR^neg^ Mϕs in the PV lacked the expression of CD163, a marker reported to be expressed in all Hofbauer cells (Reyes and Golos, 2018; Schliefsteiner et al., 2017). It is possible that CD163^lo^ Mϕs reflect downregulation of CD163 by Hofbauer cells during cell isolation as has been reported in the presence of collagenase (Tang et al., 2011). However, it is also possible that Hofbauer cell populations are more diverse than previously thought and CD163 should be used in combination with other Mϕs markers such as CD14, CD68 and DC-SIGN (Yang et al., 2017) in future studies. It is also possible that some of these cells represent non-Hofbauer macrophages within the PV.

In addition to reporting novel immune cell phenotypes, we identified mechanisms of immunosuppression and nonclassical function for multiple subsets of PV immune cells likely involved in preventing inflammation *in utero.* Specifically, we found that innate non-APCs expressed lower levels of multiple chemokine receptors including: CXCR3, CCR6, CCR4 and CCR7 than decidual counterparts. This finding is consistent with histologic evaluation of Hofbauer cells showing a lack of CCR7 and CX3CR1 staining (Joerink et al., 2011). This coupled with reduced expression of multiple chemokine ligands for these and other chemokine receptors we observed in the bulk RNAseq data, could suggest that innate cells in the PV are either more static or are mobile in a non-targeted manner at baseline.

We also report a diversity of HLA-DR^pos^ cells present in the PV, where we identified mDCs, pDCs, B cells, Mϕs and a population of HLA-DR^pos^ NK cells. An antigen-presenting role for NK cells has been previously described (Roncarolo et al., 1991). The identification of fetal HLA-DR^pos^ Mϕs contrasts Thomas et *al’s* recent findings showing no HLA-DR^pos^ cells in the PV core up to the 10^th^ week of gestation (Thomas et al., 2021). It should be noted that some of these HLA-DR^pos^ Mϕs in our study may reflect the presence of contaminate maternal Mϕs (termed PAMMs) repairing breaks in the trophoblast layer. However, the location of HLA-DR^pos^ cells in our study by both immunofluorescent and IMC show that some cells are located distant from the trophoblast layer and suggest that HLA-DR^pos^ Mϕs appear in the stroma after the time period studied by Thomas et al or between 10-18 weeks’ gestation. It would be interesting to determine the fetal versus maternal origin of these second trimester HLA-DR^pos^ Mϕs and evaluate if the Hofbauer cell population from 18-23 weeks is transcriptionally distinct from those detected in prior studies (Vento-Tormo et al., 2018).

Similar to other innate population in the PV, we detected novel immunosuppressive mechanisms in APC subsets observed within the PV. Irrespective of PV APC ontogeny, we determined that PV APCs express more PD-L1 per cell than decidual counterparts. PD-L1’s function as a coinhibitory molecule has been extensively studied (Sun et al., 2018). Moreover, PD-L1 expression can be mediated through interferon (IFNγ) signaling (Garcia-Diaz et al., 2017). Interestingly we showed that PV immune cells produce IFNγ preferentially to TNFa, another proinflammatory cytokine. These findings insinuate that PV APCs mediate in utero homeostasis by controlling T cell activation through the expression of coinhibitory ligands. Further validating the possibility of PV APC mediation of homeostasis via PD-L1 expression, we found that preterm PV have significantly reduced expression of PD-L1 on PV stromal cells compared to mid-gestation PV. Though it is not possible to determine if PD-L1 reduction precedes or is the consequence of preterm delivery, we concurrently observed increased pZAP70 in preterm PV over mid-gestation PV, consistent with increased T cell activation. Thus, it is possible that loss of PD-L1 on PV APCs results in elevated T cell activation leading to increased inflammation that is a well-documented phenotype in preterm deliveries (Romero et al., 2006). With the recent advancements in immunotherapy in multiple disease contexts it would be very interesting to study the local effects in the placenta and pregnancy outcomes of check-point blockade therapies throughout gestation.

The detection of memory T cells within the PV justifies the need for PV APCs to limit T cell activation. Memory T cells have been detected in multiple fetal human organs (Angelo et al., 2019; Halkias et al., 2019; Li et al., 2019; Schreurs et al., 2019; Stras et al., 2019b), and with in the non-human primate PV (Toothaker et al., 2020). Additionally activated and resting T cells have been detected in PV samples post-delivery (Pique-Regi et al., 2019) and central memory T cell can be found in human cord blood from preterm infants (Frascoli et al., 2018). Here we report with both CyTOF and flow cytometry that PV T cells are enriched for CD8 and DN T cells, consistent with previous findings in term placentas (Erbach et al., 1993; Kim et al., 2008). Moreover, we found that at baseline PV T cells express low activation signatures (CD69 and PD-1) potentially suggesting that PV T cells have been previously educated (hence the memory marker expression) but remain quiescent due to either the lack of antigens or direct inhibition from PD-L1^pos^ APCs. Moreover, we found no difference in baseline cytokine production by PV T cells between mid-gestation, preterm and term pregnancies, and increased proliferation of PV T cells compared to adult T cells in multiple organs further suggesting that PV T cell machinery and functionality is established early in pregnancy (prior to 17weeks’ gestation). Interestingly, we observed high production of GRZB (>75%) among all T cell subsets and NK cells at baseline in term PV. GRZB production in CD4 and CD8 T cell subsets has been previously observed in non-human primate PV T cells in the third trimester, however its direct implications for placental health remain unclear (Toothaker et al., 2020). Interestingly, we observed downregulation of GRZB in CD69^pos^ CD8 T cells and an increase in IFNγ in these same cells upon stimulation of the TCR pathway. One explanation for this phenomenon could be that PV T cells have altered cytokine production between TCR-independent and TCR-dependent activation pathways. We propose that TCR-independent activation, such as by cytokines allows for consistent production of GRZB by CD69^pos^ CD8 T cells at baseline. This TCR-independent high level of GRZB may allow PV T cells to act as a secondary layer of viral defense if the trophoblast layer is breached *in utero;* aide in the clearance of dying cells during cellular turnover as the placenta grows; or have a novel function specific to the placenta that remains to be elucidated. However, if PV T cells are activated through the TCR (experimentally measured with αCD3/CD28 antibodies), they respond by changing cytokine production from a cytotoxic predominance to favor IFNγ production. It is also possible then that PV T cells increase IFNγ upon TCR stimulation to increase PD-L1 on APCs to promote homeostasis and prevent *in utero* inflammation. Alternatively, IFNγ production in activated PV T cells from term placentas participates in the induction of the labor cascade that would be consistent with data showing that both IFNγ and TNFα from preterm cord blood memory T cells can induce uterine contractility (ref).

The presence of a placental and/or fetal microbiome as a source of potential antigens for PV T cells remains highly contested (Aagaard et al., 2014; de Goffau et al., 2019; Kuperman et al., 2020; Leiby et al., 2018; Mishra et al., 2021; Rackaityte et al., 2020; Theis et al., 2020). However, our group recently showed that xenobiotic metabolites including bacterial metabolites are present in the fetal intestine at 14 weeks’ gestation (Li et al., 2020). As such, it is possible that maternal bacteria derived peptides similarly cross the placenta and educate fetal T cells. Fetal T cell activation by maternal antigens has also been implicated in preterm birth where T cells from cord exposed to maternal antigens delivered on cord blood APCs showed increased proliferation and secretion of TNFα and IFNγ (Frascoli et al., 2018). As previously mentioned, our data indicates that T cells obtained from mid-gestation PV can be stimulated through the activation of the TCR signaling (αCD3/CD28 antibodies) and when exposed to decidual antigens but not to unmatched PBMC antigens. Of note, we found many more T cells upregulating CD69 compared to those secreting TNFα upon decidual stimulation. It is likely that there is a differential response to maternal antigens among PV T cells. Further validating differential response to stimulation by mid-gestation PV T cells, we observed high patient to patient variability in cytokine secretion following stimulation. This was also observed in preterm PV T cells. Yet, term PV T cells behaved in an orchestrated fashion across all patients. This suggests that prior term PV T cells responses have not been fully established, but by the end of healthy pregnancy PV T cells mature enough to have similar responses upon activation. This variation in PV T cells responses to stimulation could be attributed to either incomplete T cell development or incomplete exposure to an array of antigens. This variability could alternatively be attributed to PV T cells at term having a well-defined role whereas prior to complete gestation, the PV T cell response to stimulation is variable depending on the overall state of individual pregnancies. Collectively, these findings have identified unique functions of PV T cells and suggest that antigens could stimulate a proinflammatory PV T cell response upon crossing the placental barrier, however the multiple immunosuppressive mechanisms are in place as early as 17 weeks within the placenta to prevent inappropriate T cell activation including: limited chemotaxis of innate sensor cells and high expression of coinhibitory molecules by PV APCs.

Our study had several limitations of note the lack of genetic information to segregate fetal from maternal cells. It would be very interesting for a future cohort to definitively determine the origin of each PV immune cell subset identified in this work using dual *in situ* hybridization and immunodetection techniques. Furthermore, legal limitations prevented the collection of maternal and fetal blood to use for comparison.

The ability of fetal immune cells to execute mature functions has recently been validated in multiple cell types and organs throughout the fetus (Angelo et al., 2019; Frascoli et al., 2018; Halkias et al., 2019; McGovern et al., 2017; Schreurs et al., 2019; Stras et al., 2019a). As such the detection of immunosuppressive mechanisms to control this fetal immune response and prevent *in utero* inflammation is critical, particularly so at the point of contact and potential antigen exchange between mother and fetus. Throughout this study we have identified previously understudied immune cell populations within the mid-gestation placental villi. Moreover, we detected multiple mechanisms of immunosuppression utilized by these PV immune cells to help maintain homeostasis and prevent inflammation *in utero.* This work has implications for future studies to better understand the complex roles of fetal and maternal immune cells within the placenta and potentially contribute to a better understanding of immune tolerance in multiple disease contexts.

## Methods

### Placental Tissue Collection

Human products of conception were obtained through the University of Pittsburgh Biospecimen core after IRB approval (IRB# PRO18010491). Preterm placentas resultant from a variety of obstetric complications were obtained from the University of Pittsburgh MOMI Biobank. Term placentas were collected both at the University of Pittsburgh through the MOMI biobank and through the Yale University YURS Biobank from C-section deliveries devoid of obstetric complications (**Table 1**). Placental villi were separated using forceps under a light dissection microscope (Fisherbrand #420430PHF10) from the chorionic and amniotic membranes lining the chorionic plate (CP) and from the decidua basilis (referred to as decidua throughout manuscript) on the basal plate side of the placenta. Tissue was thoroughly washed with sterile PBS prior to cryopreservation and subsequent single cell isolation as previously described (Konnikova et al., 2018).

### RNA sequencing

Snap frozen placental tissues were shipped on dry ice to MedGenome for mRNA extraction and library preparation. RNA extractions were completed with the Qiagen All Prep Kit (#80204). cDNA synthesis was prepped with the Takara SMART-seq kit (#634894) and NexteraXT (FC-131-1024, Illumina) was used to fragment and add sequencing adaptors. Quality control was completed by MedGenome via Qubit Fluorometric Quantitation and TapeStation BioAnalyzer. Libraries were sequenced on the NovaSeq6000 for Paired End 150 base pairs for 90 million reads per sample.

### RNA sequencing analysis

FASTQ files were imported and subsequently analyzed with CLC Genomics Workbench 20.0 (https://digitalinsights.qiagen.com). Briefly, paired reads were first trimmed with a quality limit of 0.05, ambiguous limit of 2 with automated read through adapter trimming from the 3’-end with a maximum length of 150. Trimmed reads were then mapped to the homo_sapiens_sequence_hg38 reference sequence. Differential gene expression was computed in CLC Genomics with an Across groups ANOVA-like comparison. Significantly differentially expressed genes were delineated as those with a p-value <0.05, False-Discovery Rate <20% and fold-change > absolute value of 2. Heatmaps for gene expression were created with Morpheus (https://software.broadinstitute.org/morpheus).

### RISH

Formalin fixed samples were sectioned and embedded in paraffin by the Pitt Biospecimen Core. Staining was completed per manufacturer’s instructions for RNAScope^®^ multiplex V2 detection kit (ACD Bio) coupled with immunofluorescent protein staining for either Cytokeratin19 (ab52625 Abcam) at 1:250 dilution. Echo^®^ Revolve microscope at 20x was used to image sections. All images were batch processed using FIJI (Schindelin et al., 2012), and all edits were made to every pixel in an image identically across all patients per experiment. Quantification of cell populations was done using a custom pipeline in CellProfiler (McQuin et al., 2018).

### FISH

In situ hybridization for the Y chromosome was adapted from the protocol outlined in(Enninga et al., 2020). Briefly, slides were deparaffinized with a series of xylene and ethanol washes. Target retrieval was done at 95º^C^ for 10 minutes, slides were placed in 70% ethanol, 85% ethanol and 100% ethanol for 2 minutes each. DYZ3 probe (D5J10-034, Abbott Laboratories) was diluted 1:10 in LSI/WCP hybridization buffer (D6J67-011, Abbott Laboratories) and incubated for 5 minutes at 83º^C^ prior to overnight hybridization at 37º^C^. Slides were soaked in SSC/0.1% NP-40 (ab142227, Abcam) to remove cover slips and placed in 2X SSC/0.1% NP-40 for 2 minutes at 74º^C^ before mounting with antifade plus Propidium Iodide (p36935, Invitrogen). Slides were imaged on the LSM 710 (Leica Biosystems) confocal at the Yale Center for Cellular and Molecular Imaging.

### RNA extraction and qPCR

RNA was extracted from snap-frozen villi samples using the RNAEasy Minikit (#217004, Qiagen) RNA was converted to cDNA using iScript (#1708891, BioRad) reagents according to manufacturer protocol. Samples were run on the Taqman StepOnePlus Real-Time PCR System (Applied Biosciences) machine with probes for ACTB (Hs01060665_g1) as housekeeping gene and with either XIST (Hs01079824_m1) or EIF3AY (Hs01040047) all from Qiagen. Values undeterminable were given cycle values of 40 for quantification purposes.

### Immunofluorescent staining

Slides with 10um sections of FFPE tissue were deparaffinized with a series of xylene and ethanol washes. Antigen retrieval was performed in the Biocare Medical LLC decloaking chamber (NC0436641) for 1 hour with citrate-based antigen retrieval buffer (H-3300, Vector Laboratories) and washed with PBS. Slides were then blocked for 30 minutes with 10% horse serum prior to overnight incubation at 4ºC with primary antibodies. Slides were washed with PBS and incubated with secondary antibodies for 45 minutes at RT. Slides were mounted with Antifade mounting media + DAPI (H-1300, Vectashield).

### Imaging mass cytometry

Slides with 4um sections of FFPE tissue were deparaffinized with a series of xylene and ethanol washes. Antigen retrieval was performed at 95º^C^ for 20 minutes using 1X Antigen Retrieval Buffer (#CTS013 R&D) and washed with water and dPBS. Slides were then blocked for 30 minutes with 3% BSA in dPBS prior to overnight incubation at 4ºC with a primary antibody cocktail **(Table S8)**. Slides were rinsed and co-stained with 191/193 DNA-intercalator (Fluidigm), rinsed and air dried for >20 minutes prior to analysis. Slides were analyzed on the Hyperion Mass Cytometer with an ablation energy of 4 and frequency of 100Hz for ~30 minutes per section. Representative images were generated using Histocat++ software(Catena et al., 2018).

### CyTOF staining

Samples were stained with antibody cocktail (**Table S4**) per previously published protocol(Stras et al., 2019a) and incubated with 191Ir/193Ir DNA intercalator (Fluidigm) and shipped overnight to the Longwood Medical Area CyTOF Core. Data was normalized and exported as FCS files, downloaded and uploaded to Premium Cytobank^®^ platform. Any files with insufficient cell number were excluded from analysis **(Table S5)**. Gating and analysis was completed with cytofkit(Chen et al., 2016) as published (Stras et al., 2019a). Cluster abundance was extracted, and statistically analyzed using R.

### Stimulation of PV T cells

*αCD3/αCD28 with CFSE:* Cells were isolated from cryopreserved PV samples as described throughout manuscript. Dead cells were removed prior to stimulation using Millitenyl dead cell removal kit (130-090-101 Millitenyl Biotec). Cells were incubated with CFSE (65-0850-85) alone or with αCD3 (clone HIT3a, #300302, Biolegend) and αCD28 (clone CD28.2, 302902, Biolegend) soluble antibodies for 72 hours rotating at 37º^C^ + 5% CO_2_. GolgiPlug (51-2301K2, BD Biosciences) and GolgiStop (51-2092K2, BD Biosciences) were added for the last 4 hours of stimulation. *αCD3/αCD28 with Ki67:* Cells were isolated as described above and incubated with aforementioned αCD3 and αCD28 antibodies for 4 hours rotating at 37º^C^ + 5% CO_2_.

### Stimulation of PV T cells lysed decidual cells

Single cells from PV and decidua were isolated from cryopreserved tissue as previously described(Konnikova et al., 2018). PBMCs were thawed and DMSO was washed out. PV tissue was thawed and made into single-cell suspensions (as described above). Cells were incubated in 5mLs of media with GolgiPlug (51-2301K2, BD Biosciences) and GolgiStop (51-2092K2, BD Biosciences) and designated stimuli. For PMA condition, PMA (1:2000) (Sigma-Aldrich) and Ionomycin (1:1000) (Sigma-Aldrich) were added. Decidua and PBMC cells were lysed via ultracentrifugation at max speed for 7 minutes and 1mL of lysed components was added to appropriate conditions. PV cells were exposed to stimuli for 4 hours at 37º^C^ with 5% CO_2_.

### Flow cytometry

Post stimulation cells were washed with PBS and incubated with either Propidium Iodide or Zombie Aqua live/dead stain. Viability marker was washed out and cells were resuspended and spun down in FACS buffer then incubated with Human TruStain FcX (Biolegend) for 10 minutes prior to the addition or a surface antibody cocktail **(Table S9)** Cells were washed with FACs buffer and permeabilized with FoxP3 fix/perm (Invitrogen) overnight. Cells were washed with 1X FoxP3 Wash Buffer (Invitrogen) and incubated with intracellular antibodies÷Cells were washed, fixed for 10 minutes with 4% PFA and resuspended in FACs buffer. All samples were run either on All samples were run on BD LSRFortessa (BDBiosciences) at the University of Pittsburgh Department of Pediatrics Flow Cytometry core (decidual cell stimulation) or on BD LSRII (BDBiosciences) at the Yale University Flow Cytometry core (all other stimulations).Output FCS files were analyzed with FlowJo ^®^.

### Statistics

R version 3.6.1 with Kruskal-Wallis analysis and Dunn’s multiple comparison test for post-hoc analysis among groups. One-tailed t-test was used to compare groups of two. Comparisons of mean expression values corrected using the Bonferroni method. P-values of 0.05 or less were significant.

### Plot generation

Plots comparing multiple groups were generated using Prism GraphPad 8. In each plot, each data point represents one subject as per figure description.

## Data and code availability

Data analyzed in this study has been stored according to IRB guidelines and is subject to institutional regulations. Requests can be directed to Lead Contact, Liza Konnikova.

## Acknowledgements

This project used the UPMC Hillman Cancer Center and Tissue and Research Pathology/Pitt Biospecimen Core shared resource which is supported in part by award P30CA047904. This research was supported in part by the University of Pittsburgh Center for Research Computing through the resources provided. We thank Yale Flow Cytometry for their assistance with LSRII service. The Core is supported in part by an NCI Cancer Center Support Grant # NIH P30 CA016359. We also thank Ansen Burr and the Hand Lab at The University of Pittsburgh for their help with RNA-sequencing analysis and Meghan Mooring and Dean Yimlamai for help with confocal microscopy and Biorender images.

## Author Contributions

JMT and LK conceived the work. CyTOF and flow cytometry procedures and analysis and drafting of manuscript were done by JMT supervised by LK. Tissue collection and preparation was done by JMT, RMC, CCM and OOO. IMC analysis was done by OOO. Immunocytochemistry was completed by BTM; RISH was done by RMC; RNA extraction/qPCR was performed by CCM. Cytometry bioinformatics consultation was done with PL supervised by GT. RNA sequencing assistance provided by DY. Figure construction was done by JMT and BTM. All authors contributed to the editing and compilation of the manuscript.

## Supplemental Tables included at end of document

Table S1. Patient Cohort

Table S2. Differentially Expressed Genes Between Decidua, CP and PV

Table S3. Selected Immune Genes

Table S4. CyTOF Panel

Table S5. Files Omitted from CyTOF Analyses

Table S6. Cell Type Identification

Table S7. IMC Panel

Table S8. Differentially Expressed Chemokines

Table S9. Flow Cytometry Antibodies

---

**Figure S1.**
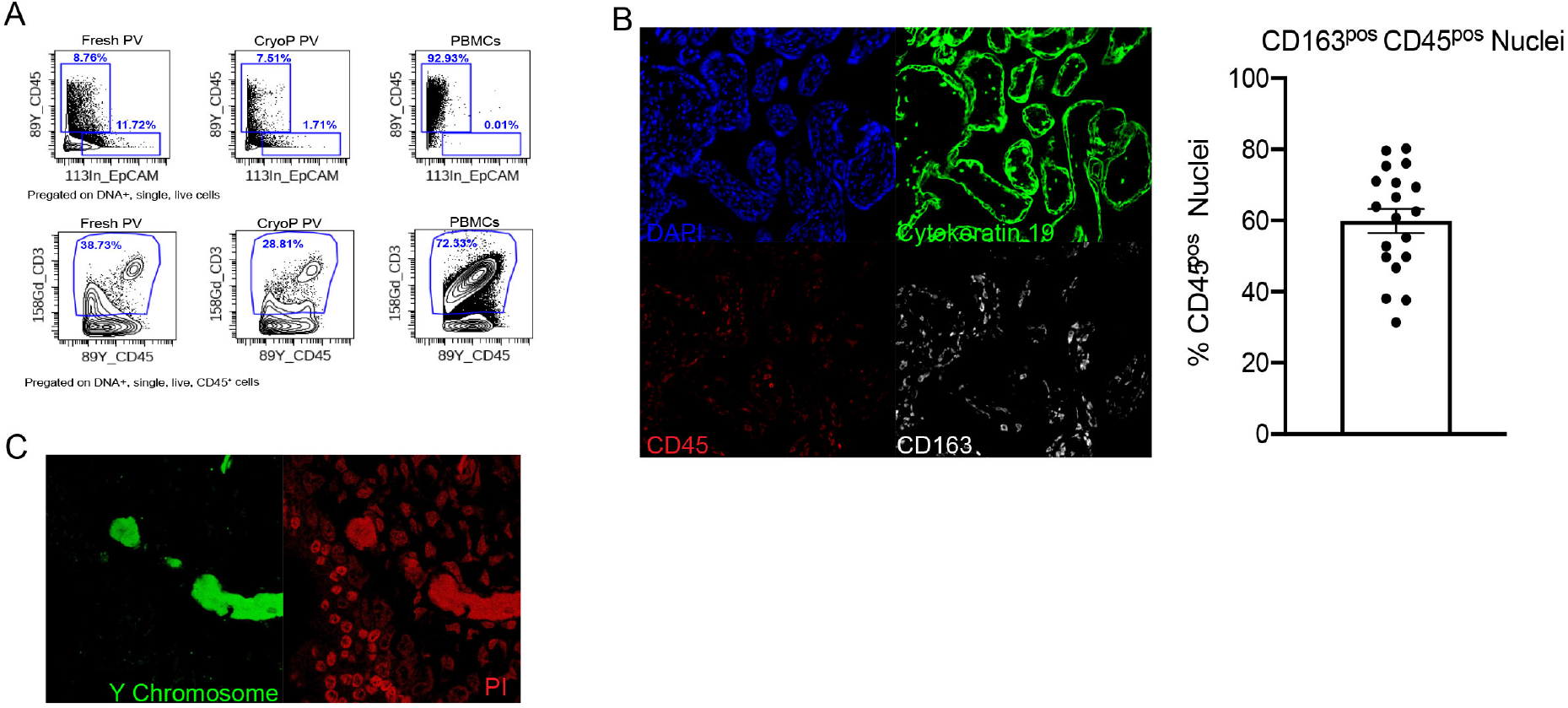
(A) Comparison of CyTOF analysis of CD45^pos^ and T cell abundances between fresh PV, cryopreserved PV and frozen Peripheral Blood Mononuclear Cells (PBMCs). (B) Splits of immunofluorescence for intravillous immune cell detection and quantification of CD163^hi^ immune cells. (C) Splits for Y-chromosome *in situ* hybridization.

**Figure S2.**
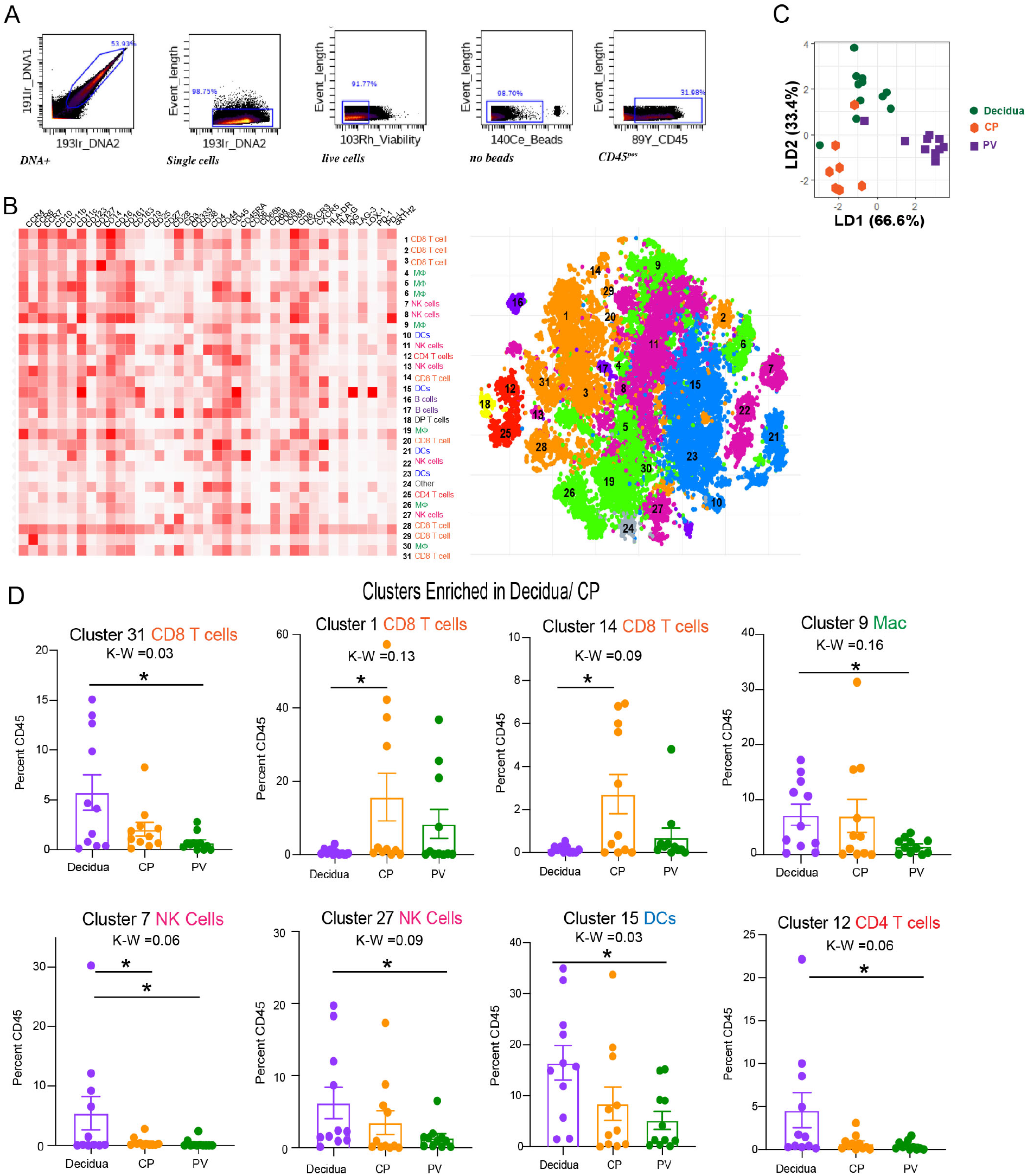
(A) Pre-gating strategy for CD45^pos^ population used in automated clustering. (B) Clustergrammar heatmap used for cluster identification (left) of tsne clusters mapped (right). (C) Linkage Disequilibrium (LD) plot confirming segregation of placental layer immune profile. (D) Abundance of decidua and CP enriched clusters. * = p value < 0.05 after posthoc analysis from Kruskal-Wallis (K-W) test.

**Figure S3.**
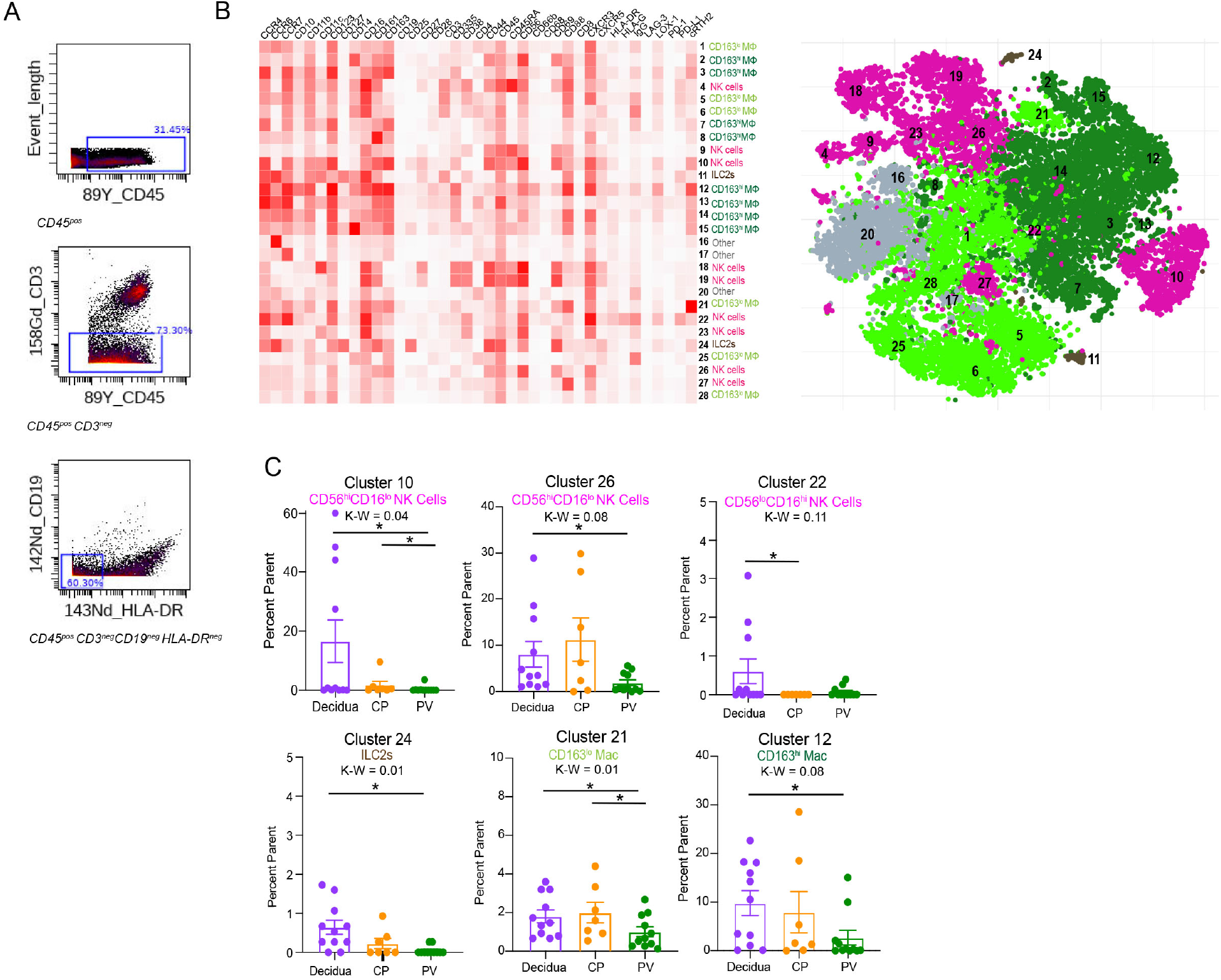
(A) Pregating strategy for innate non-APC population. (B) Clustergrammar heatmap used for cluster identification (left) of tsne clusters mapped (right). (C) Cumulative data on abundance of decidua and CP enriched clusters. * = p value < 0.05 after posthoc analysis from Kruskal-Wallis (K-W) test.

**Figure S4.**
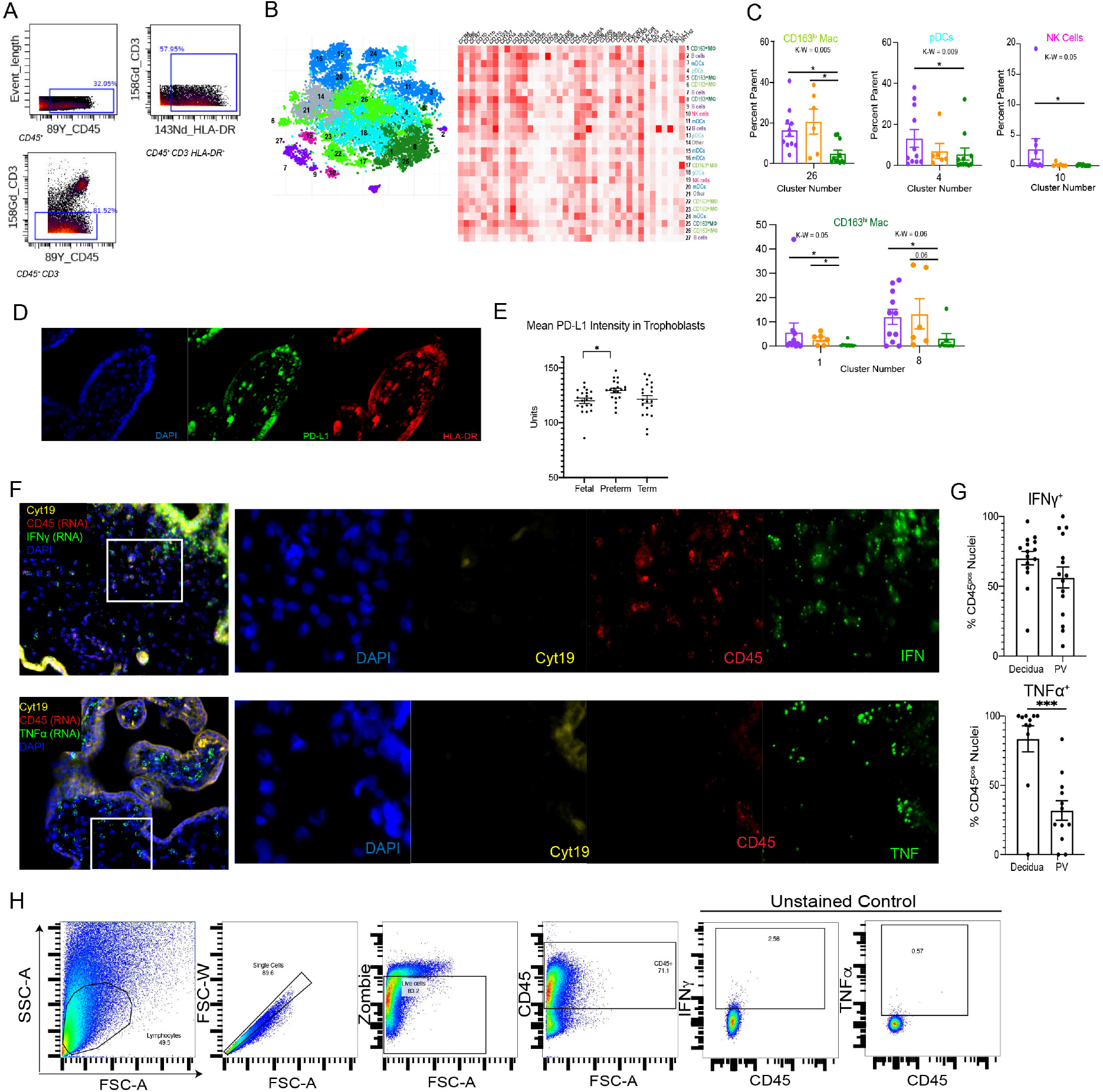
(A) Pregating strategy for APC population. (B) Clustergrammar heatmap used for cluster identification (left) of tsne clusters mapped (right). (C) Cumulative data on abundance of decidua and CP enriched clusters. * = p value < 0.05 after posthoc analysis from Kruskal-Wallis (K-W) test. (D) Splits for PD-L1^pos^ PV APCs representative image. (E) Quantification of trophoblast expression of PD-L1 via IHC. (F) Splits for dual RNA *in situ* hybridization and IF. Representative images in main figure taken from region identified in white rectangle. (F) Quantification of staining for cytokine positive immune cells in PV and decidua. (H) Pregating for CD45^pos^ population with flow cytometry. *** = p-value < 0.001 in Mann-Whitney two tailed test.

**Figure S5.**
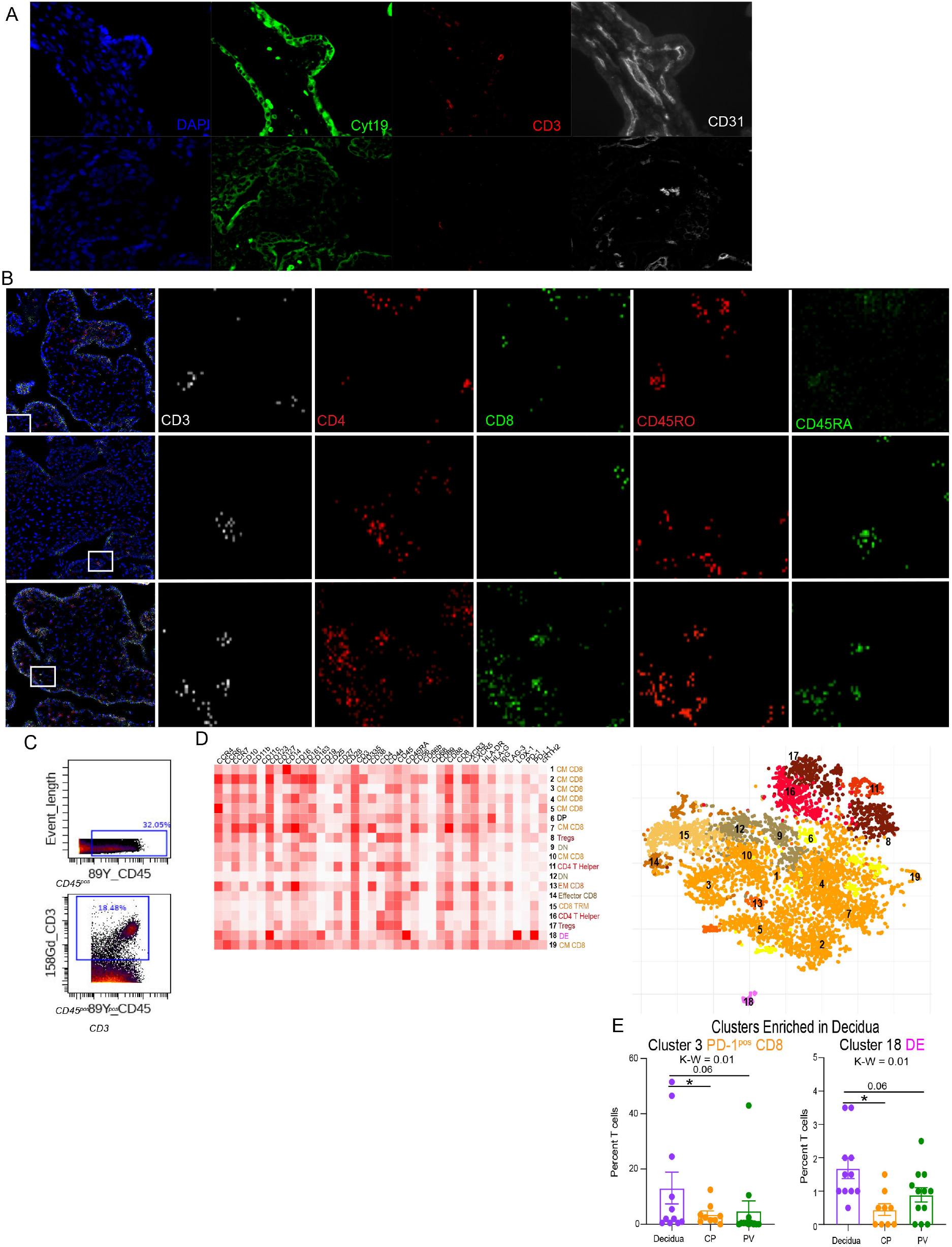
(A) Splits from IF detection of T cells and PV endothelium. (B) Splits from representative IMC images of T cell subsets. T cells show in main figures identified in regions highlighted with white rectangles. (C) Pregating strategy for T cell population. (D) Clustergrammer heatmap used for cluster identification (left) of tsne clusters mapped (right). (E) Cumulative data on abundance of decidua and CP enriched clusters. * = p value < 0.05 after posthoc analysis from Kruskal-Wallis (K-W) test.

**Figure S6.**
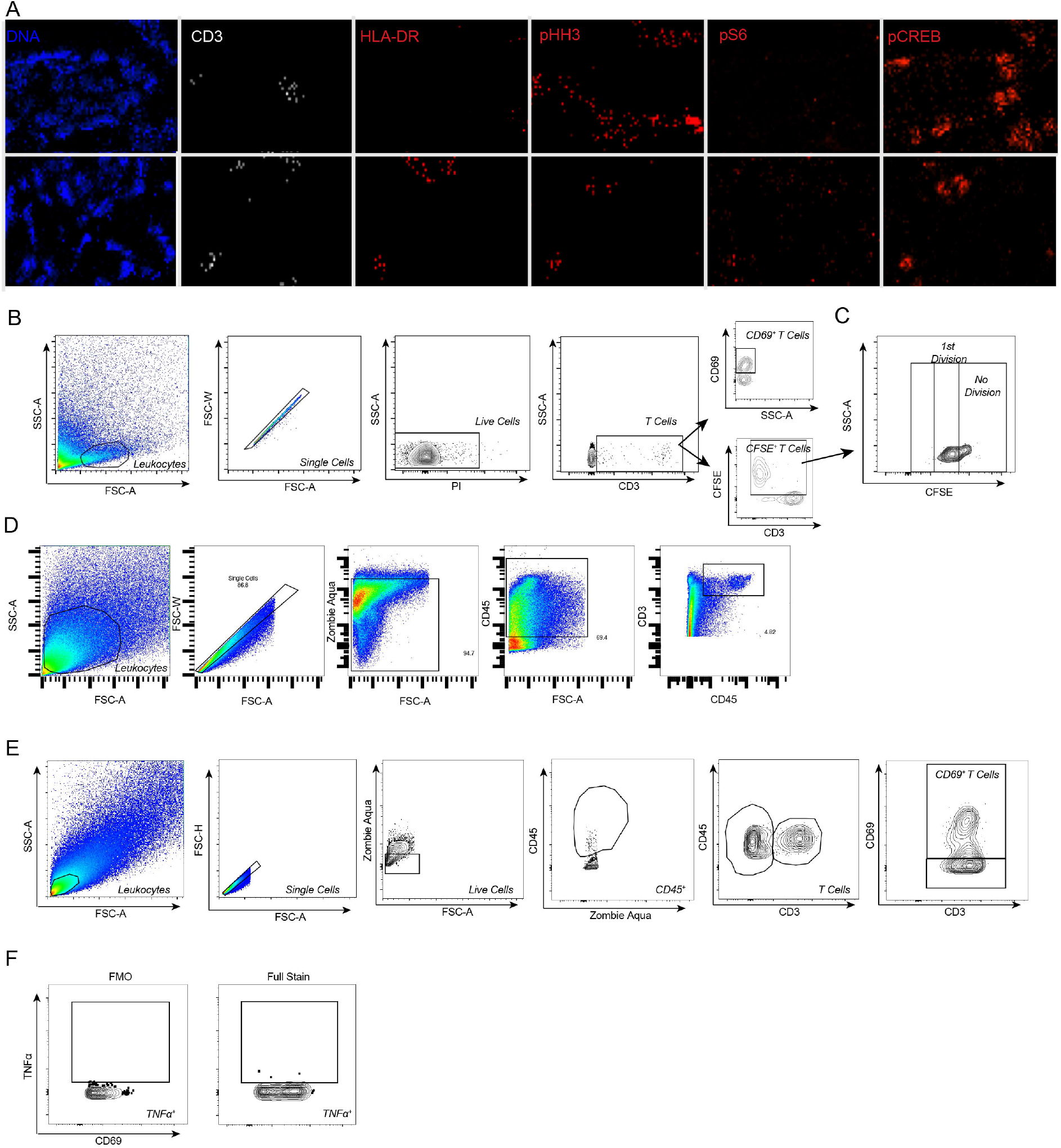
(A) Splits from IMC images showing inactive and active T cells in PV. (B) Pregating strategy for CD69^pos^ and CFSE^pos^ T cells. (C) Gating for proliferative populations quantified in main text. (D) Gating for T cells in Ki67 proliferation experiment in main text. (E) Pregating strategy for CD69^pos^ T cells used in decidual stimulation. (F) Gating of TNFα^pos^ population.

**Figure S7.**
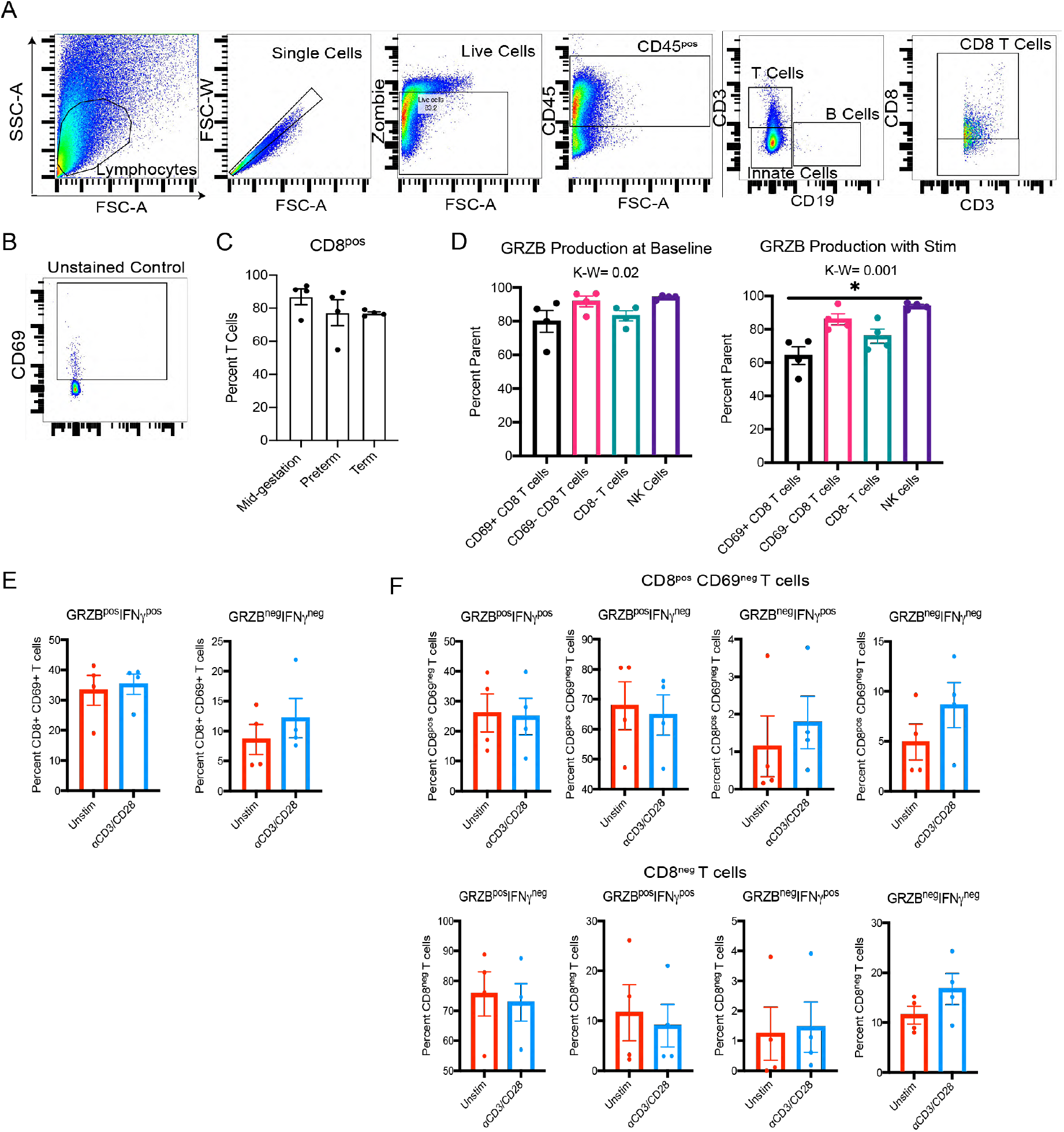
(A) Pregating of flow cytometry to CD8 T cells. (B) Unstained control for CD69 and cytokine stains in flow cytometry. (C) Quantification of CD8 T cell abundance in flow cytometry. (D) Quantification of GRZB by T cell and NK subsets from flow cytometry. (E) Quantification of GRZB and IFNγ populations in CD8^pos^ CD69^pos^ T cells. (F) Quantification of GRZB and IFNγ populations in T cell subsets.

